# Estrogen-independent molecular actions of mutant estrogen receptor alpha in endometrial cancer

**DOI:** 10.1101/369165

**Authors:** Zannel Blanchard, Jeffery M. Vahrenkamp, Kristofer C. Berrett, Spencer Arnesen, Jason Gertz

## Abstract

Estrogen receptor 1 (*ESR1*) mutations have been identified in hormone therapy resistant breast cancer and primary endometrial cancer. Analyses in breast cancer suggests that mutant ESR1 exhibits estrogen independent activity. In endometrial cancer, *ESR1* mutations are associated with worse outcomes and less obesity, however experimental investigation of these mutations has not been performed. Using a unique CRISPR/Cas9 strategy, we introduced the D538G mutation, a common endometrial cancer mutation that alters the ligand binding domain of ESR1, while epitope tagging the endogenous locus. We discovered estrogen-independent mutant ESR1 genomic binding that is significantly altered from wildtype ESR1. The D538G mutation impacted expression, including a large set of non-estrogen regulated genes, and chromatin accessibility, with most affected loci bound by mutant ESR1. Mutant ESR1 is unique from constitutive ESR1 activity as mutant-specific changes are not recapitulated with prolonged estrogen exposure. Overall, D538G mutant *ESR1* confers estrogen-independent activity while causing additional regulatory changes in endometrial cancer cells that are distinct from breast cancer cells.

## Introduction

Estrogen receptor 1 (ESR1) is a ligand-inducible steroid hormone receptor that acts as an oncogene in many breast and endometrial tumors. In these diseases, hormone therapies can be used to reduce estrogen signaling either through a reduction in estrogen production or a reduction in ESR1 activity. Mutations in the ligand binding domain (LBD) of *ESR1* have been associated with hormone therapy resistance in breast cancer (Fuqua 1993; Osbourne 2011; Robinson et al. 2013; Toy et al. 2013; Fuqua et al. 2014; Jeselsohn et al. 2014; Toy et al. 2017) and a recent large-scale analysis of *ESR1* mutations found that 14% of metastatic breast cancers harbor a LBD mutation (Toy et al. 2017). *ESR1* LBD mutations were not identified in primary tumors in The Cancer Genome Atlas’ study on breast cancer (Network 2012), indicating that *ESR1* mutations are not observed at near clonal frequencies and are unlikely to play a role in tumor initiation; however, *ESR1* LBD mutations can be found at low mutation frequencies in primary breast tumors (Toy et al. 2017). In contrast, heterozygous *ESR1* LBD mutations are found in 5.8% of primary endometrial cancers with endometrioid histology (Cancer Genome Atlas Research et al. 2013; Backes et al. 2016; Gibson et al. 2016), representing approximately 1,500 new uterine cancer diagnoses with an *ESR1* LBD mutation in the United States each year. The presence of *ESR1* mutations is associated with obesity-independent endometrial cancer and patients with *ESR1* LBD mutations trend towards worse prognosis when compared to patients with wildtype *ESR1* tumors (Backes et al. 2016).

The *ESR1* LBD mutations occur in a region of the protein essential for ligand binding and interactions with co-regulatory proteins, with the majority of mutations found at residues D538 and Y537. Studies into the molecular and phenotypic consequences of *ESR1* LBD mutations have been performed in breast cancer, revealing that the mutations confer estrogen-independent ESR1 activity, which drives gene regulation and cell proliferation in the absence of estrogens (Merenbakh-Lamin et al. 2013; Robinson et al. 2013; Toy et al. 2013; Jeselsohn et al. 2014; Bahreini 2017; Toy et al. 2017; Zhao 2017; Jeselsohn 2018). Biochemical characterization of the mutations suggests that mutant ESR1 favors the activated conformation of the receptor irrespective of ligand, causing constitutive receptor activity (Merenbakh-Lamin et al. 2013; Fanning 2016; Toy et al. 2017; Zhao 2017; Katzenellenbogen 2018). Gene expression analyses highlight the ability of mutant ESR1 to regulate canonical ESR1 target genes in the absence of estrogens (Merenbakh-Lamin et al. 2013; Robinson et al. 2013; Toy et al. 2013; Jeselsohn et al. 2014; Bahreini 2017; Toy et al. 2017; Jeselsohn 2018; Katzenellenbogen 2018). In addition to ligand-independent regulation of genes that are normally impacted by 17β-estradiol (E2), novel non-E2 regulated genes are also affected by mutant *ESR1* (Bahreini 2017; Jeselsohn 2018), suggesting that these mutations may confer additional functionality to ESR1 than just constitutive activity. While these studies have uncovered important features of the molecular and phenotypic consequences of *ESR1* mutations in breast cancer, similar analyses have not been performed in endometrial cancer cells. Because gene expression responses to estrogens and ESR1 genomic binding are highly dissimilar between breast and endometrial cancer (Gertz 2012; Gertz et al. 2013; Droog 2017), the impact of *ESR1* LBD mutations in endometrial cancer cells could be different than the effects observed in breast cancer cells.

In this study, we sought to gain an understanding of the molecular consequences of *ESR1* LBD mutations in endometrial cancer. Utilizing Ishikawa cells, a human endometrial adenocarcinoma cell line that is a cell culture model for type I disease, we employed a CRISPR/Cas9 mediated epitope tagging strategy. We created endometrial cancer cells that are heterozygous for the D538G *ESR1* ligand binding domain mutation (or wildtype *ESR1* for controls) coupled with a FLAG epitope tag incorporated at the C-terminus of the endogenous locus. The addition of an epitope tag to the endogenous gene allowed us to specifically analyze binding of the mutant form of ESR1 by Chromatin immunoprecipitation followed by high-throughput sequencing (ChIP-seq). We explored mutation-specific gene expression effects via RNA-seq, assessed changes to the chromatin landscape using the Assay for Transposase Accessible Chromatin followed by sequencing (ATAC-seq), and analyzed effects on proliferation and migration. We also investigated whether the regulatory effects of mutant ESR1 could be recapitulated by prolonged exposure to E2. The systematic investigation of mutant ESR1’s molecular activity in endometrial cancer cells will enable future phenotypic and mechanistic investigation into *ESR1* mutant endometrial cancer.

## Results

### Generation of D538G *ESR1* mutant and wildtype cell lines

The molecular consequences of *ESR1* LBD mutations have not been explored in endometrial cancer and warrant investigation. We utilized a CRISPR/Cas9-mediated epitope tagging strategy called CETCH-seq (Savic et al. 2015) in Ishikawa cells, an endometrial adenocarcinoma cell line that exhibits ESR1 genomic binding similar to endometrial tumor samples (Rodriguez et al. 2019), to model a common *ESR1* LBD mutation, D538G. This technique combines guide RNAs that target Cas9 to the C-terminus of ESR1 and a donor plasmid that leads to the incorporation of a 3X FLAG epitope tag and neomycin resistance gene at *ESR1*’s endogenous locus (Figure 1A). Ishikawa cells were first transfected with plasmids, treated with G418 to select for resistant cells and subjected to limiting dilution plating to generate single cell clones. Using this technique, we generated three cell lines that were heterozygous for the D538G *ESR1* mutation on a FLAG-tagged allele and two heterozygous FLAG-tagged wildtype cell lines (referred to as wildtype clones throughout, the original Ishikawa cell line is referred to as parental). FLAG and ESR1 protein expression were established via western blot (Figure 1B) and D538G mutations were confirmed by Sanger sequencing. Additionally, we observed similar expression frequencies of the wildtype and mutant alleles in the D538G clonal cell lines by RNA sequencing (RNA-seq) (Figure 1C). The creation of these isogenic cell lines enabled further studies into the gene regulatory changes caused by the D538G *ESR1* mutation.

**Figure 1.**
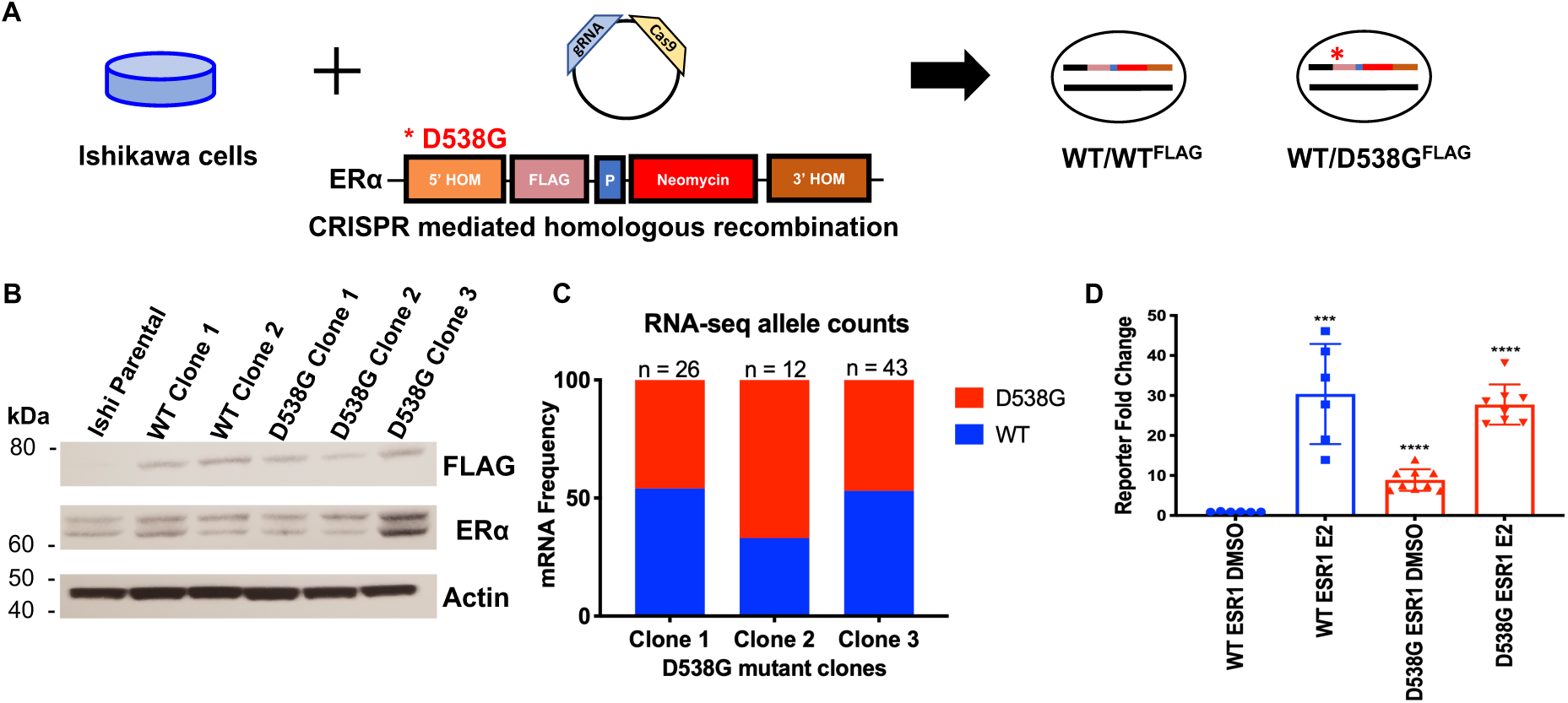
Generation and characterization of ESR1 wildtype and D538G mutant models. **A.** CRISPR-mediated epitope tagging strategy was used to generate heterozygous FLAG-tagged wildtype ER and D538G mutant ER Ishikawa cell lines. **B.** Immunoblotting for FLAG and ER in Ishikawa parental cells, two heterozygous FLAG-tagged wildtype and three heterozygous FLAG-tagged D538G mutant cell lines show protein expression of epitope-tagged ER and total ER. **C.** The ER wildtype and mutant allele expression frequencies based on RNA-seq data is shown for each D538G clonal cell line. **D.** ERE reporter activity as measured by luciferase activity was assayed in DMSO and E2 induced conditions. Experiments were done in triplicate and the figure shows the average luciferase activity for two wildtype and three D538G mutant clones. (*** p = 0.0002, ****p < 0.0001). Error bars represent SEM.

### D538G mutant ESR1 displays ligand-independent regulatory activity

To assess ESR1’s transcriptional activity in our models, we cultured wildtype and D538G mutant cells in hormone deprived media for 5 days and then transfected the cell lines with a luciferase estrogen response element (ERE) reporter assay. One day post transfection, cells were treated with either vehicle (DMSO) or 10 nM E2 for 24 hours before measuring luciferase activity. We observed negligible luciferase expression in the wildtype cells treated with DMSO (Figure 1D). However, there was a significant 30-fold increase in expression in wildtype cells induced with E2. We detected ESR1 activity in the D538G mutant clones treated with DMSO at 30% of the wildtype E2 levels. Activity was significantly increased in the D538G mutant lines treated with E2, with levels similar to the wildtype lines treated with E2. These results indicate ligand-independent transcriptional activity of the D538G mutant in endometrial cancer cell lines.

### Introduction of the D538G *ESR1* mutation causes large transcriptional changes

The ligand-independent transcriptional activity of mutant ESR1 in the reporter assay suggested that transcription in these mutant lines could be altered, leading to aberrant gene expression. To uncover gene expression changes caused by the mutation, we performed RNA-seq on the wildtype FLAG-tagged clones and D538G mutant clones following an 8 hour 10 nM E2 induction in hormone deprived media. Principal component analysis of the isogenic lines clustered wildtype and mutant lines separately, with the first principal component accounting for 46% of the variance in our datasets while separating samples based on the presence of the mutation (Figure S1A). Treatment with E2 along with differences between clones with the same genotype accounted for 29% of the variance in these lines and is represented in the second principal component, highlighting the importance of analyzing multiple clones in order to capture the variance in clone derivation. Our analysis indicates that the gene expression profile of wildtype cells supplemented with E2 did not recapitulate the expression changes seen in the D538G mutant lines.

Analysis of differentially expressed genes across clonal cell lines revealed multiple expression patterns (Figure 2A, see Table S1 for gene lists). We identified 119 genes that were upregulated and 48 genes that were downregulated in response to an estrogen induction in wildtype cells (adjusted p-value < 0.05). A comparison of these E2 responsive genes to genes regulated in the D538G DMSO treated lines, revealed 47 upregulated and 10 downregulated genes that were responsive to estrogen in wildtype cells and also regulated by the D538G mutation independent of E2. Examples of mutant ESR1 ligand-independent regulation of estrogen responsive genes confirmed by qPCR include progesterone receptor (*PGR*) known to play critical roles in reproductive function (Figure 2B) and a matrix metallopeptidase (*MMP17*) which has been implicated in the degradation of the extracellular matrix (Figure 2C). The estrogen-independent regulation of these genes by mutant ESR1 is consistent with the reporter assays and the hypothesis that mutant ESR1 has ligand-independent activity.

**Figure 2.**
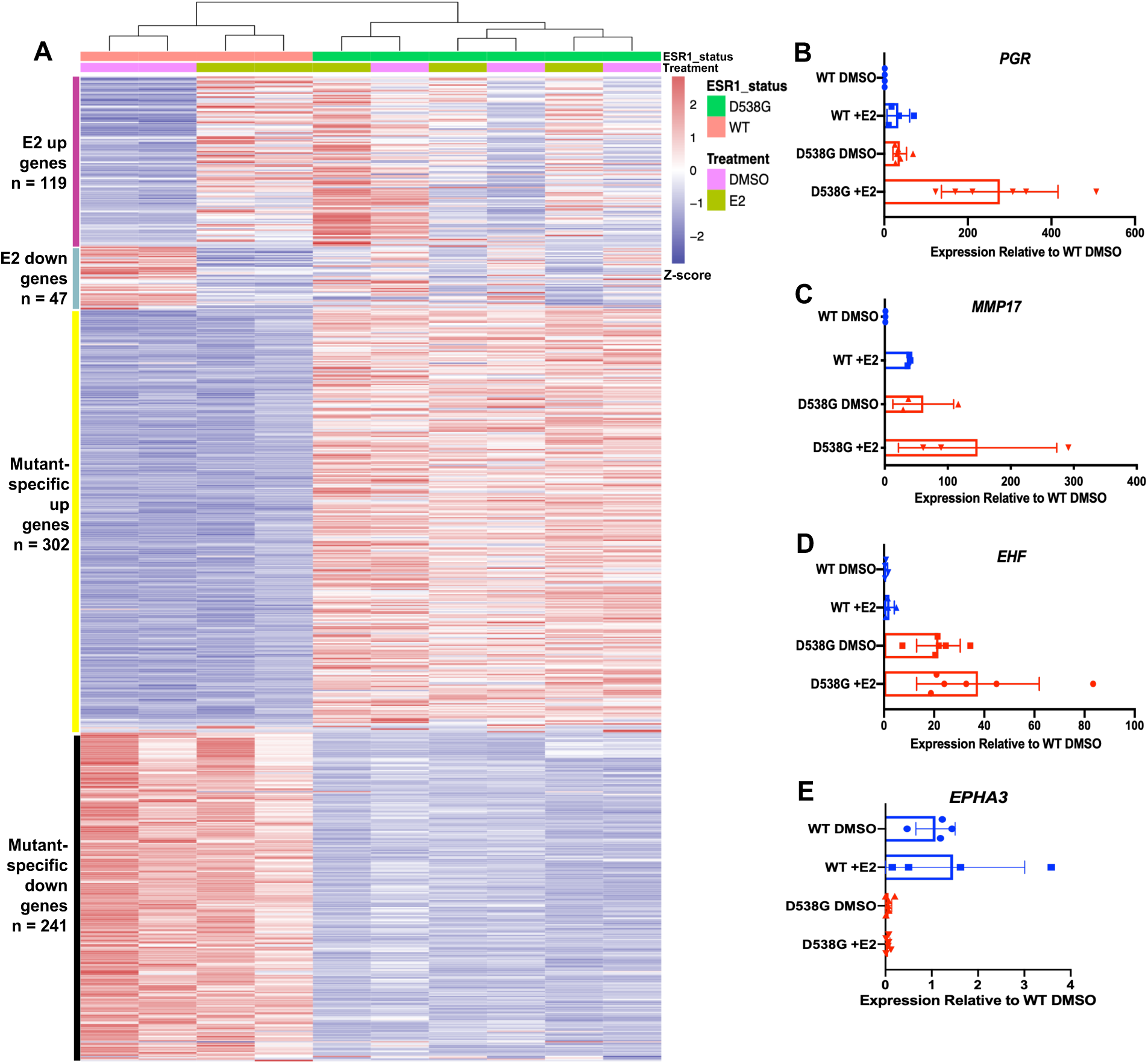
D538G mutant drives a distinct transcriptional program. **A.** Heatmap shows the relative expression of E2 up- and down-regulated genes as well as mutant-specific differentially expressed genes. Sample types are indicated by the column annotations described in the legend. Validation of RNA-seq results by qPCR are shown for ligand-independent E2 upregulated genes PGR **(B)** and MMP17 **(C)** as well as mutant-specific upregulated gene EHF **(D)** and downregulated gene EPHA3 **(E)**. Figures B-E show average expression levels, normalized to wildtype cells without E2 treatment, for two ESR1 wildtype and three D538G mutant clones after 8 hour E2 or DMSO (vehicle) inductions. Error bars represent SEM.

In addition to changes in estrogen regulated genes, the D538G mutation impacted the expression of many genes not normally regulated by estrogen. We identified mutation-specific gene expression changes with 302 novel genes that are upregulated and 241 novel genes that are downregulated due to the mutation (see Table S1 for mutant-specific genes). Examples of novel genes include *EHF*, an ETS factor not normally expressed in Ishikawa cells (Figure 2D) and EPHA3, a receptor tyrosine kinase (Figure 2E). Ingenuity Pathway Analysis indicated that novel genes were enriched for pathways associated with more aggressive tumors, which include cellular growth, proliferation and movement (Figure S1B,C). This pathway enrichment is similar to findings from studies done on *ESR1* mutations in MCF-7 and T47-D, two breast cancer cell lines. While the pathways are consistent with novel gene regulation observed in breast cancer cells, there is little overlap in the differentially expressed genes from the Bahreini et al study (MCF-7: 29 genes (5.3%); T-47D 20 genes (3.7%))(Bahreini 2017) and the Jeselsohn et al study (MCF-7: 91 genes (16.7%))(Jeselsohn 2018). Collectively, our data indicates that the D538G mutation impacts many non-E2 regulated genes and enables a more expansive and potentially aggressive transcription program.

While there are not enough *ESR1* mutant patient samples with associated gene expression data available to determine if the observed mutant-specific gene expression changes are seen in endometrial tumors, we can determine if the expression of these mutant-specific genes are associated with patient outcomes. To explore this connection, we analyzed gene expression and disease-free survival in the TCGA endometrial cancer cohort (Cancer Genome Atlas Research et al. 2013) and restricted the analysis to high *ESR1* expressing tumors with endometrioid histology. We found significant overlaps between mutant regulated genes and genes whose expression is associated with disease-free survival (see Figure S2 for examples and Table S2 for the full gene list). For the mutant up-regulated genes, there was 3.2-fold enrichment specifically for genes associated with worse outcomes (p-value = 2.7×10^-4^, Fisher’s exact test, Figure S2). And the mutant down-regulated genes were enriched 2.8-fold over random chance specifically in genes whose expression was associated with longer disease-free survival (p-value = 6.9×10^-3^, Fisher’s exact test, Figure S2). These results suggest that genes regulated by mutant ESR1, that are unrelated to E2 inductions, tend to be associated with outcomes in endometrial cancer patients in a pattern consistent with mutant ESR1 driving more aggressive tumors.

### D538G mutation increases migration without altering proliferation

The cellular and molecular pathways found enriched in D538G mutant-specific gene lists suggested a more aggressive phenotype in endometrial cancer cells. To determine how the mutation affected proliferation, wildtype and D538G mutant cells were seeded at low densities in full serum media and hormone deprived media for up to 72 hours. Growth rates were measured on the IncuCyte Zoom live-cell imaging platform. The D538G mutation did not enhance proliferation in full serum media when compared to *ESR1* wildtype cells (Figure 3A), with all cell lines showing similar growth rates. In hormone deprived media, although we observed differences in growth rates between clones, there were not consistent differences between *ESR1* wildtype and D538G mutant lines. We also tested the D538G mutant’s ability to affect migration via a wound healing assay. *ESR1* wildtype and D538G mutant cells were initially grown to 100% confluency in full serum and hormone deprived media, the cell monolayer was then scraped and imaged for up to 24 hours on the IncuCyte Zoom. The D538G mutation enhanced migration by 40.3% compared to wildtype lines in full serum media (p-value <0.0001, unpaired *t*-test, Figures 3B,C) and by 35.4% in hormone deprived media (p-value <0.0001, unpaired *t*-test, Figure 3D,E), suggesting that the mutation confers a more migratory phenotype. These results are consistent with mutant *ESR1* contributing to more aggressive endometrial tumors.

**Figure 3.**
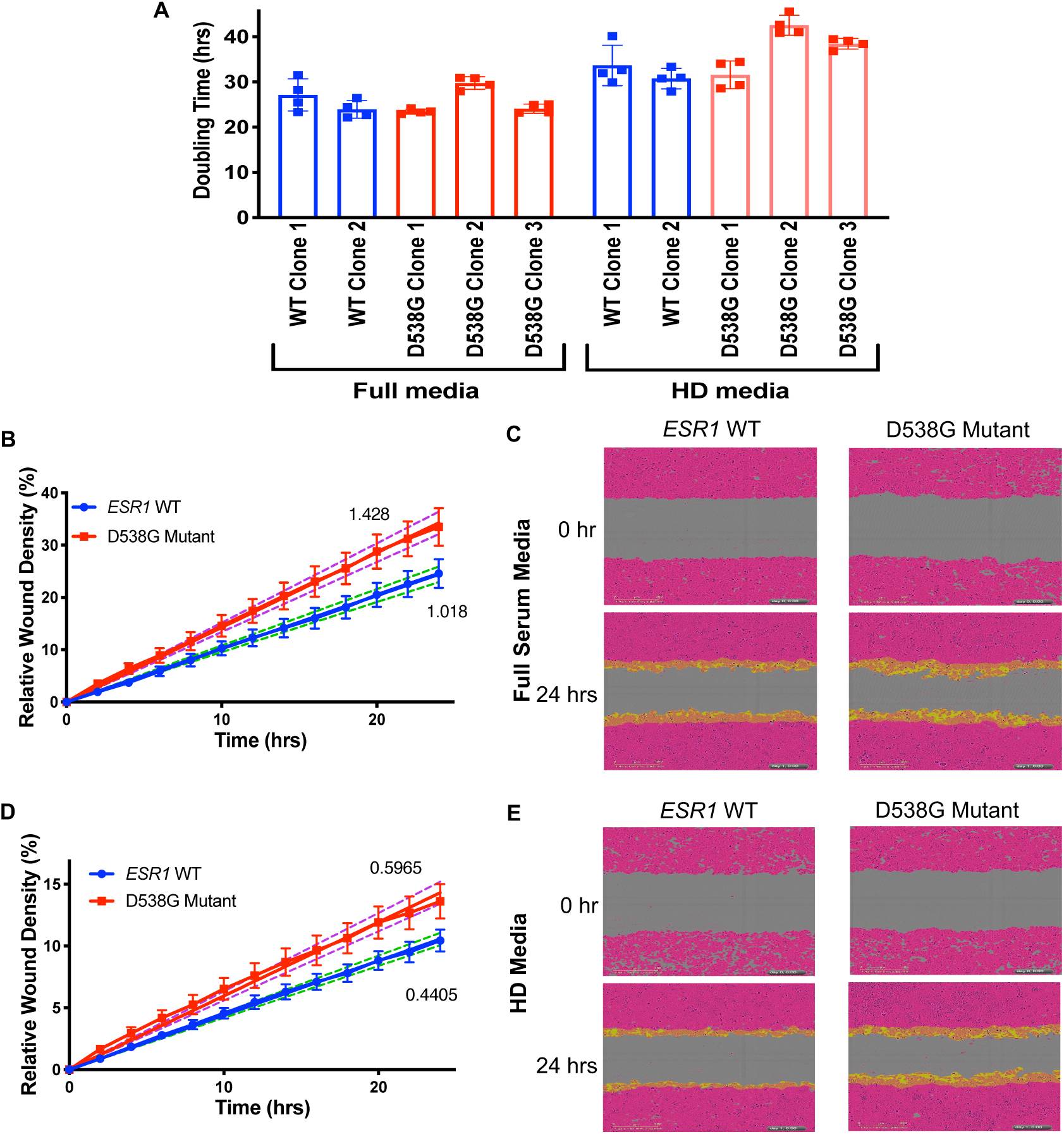
D538G mutant affects migration but not proliferation in endometrial cancer cells. **A.** Bar graphs indicating the doubling times for ESR1 wildtype and D538G mutant cell lines in full media and hormone deprived (HD) media. The relative wound densities of two ESR1 wildtype and three D538G mutant cell lines are shown over 24 hours following scratch/wounding of cell monolayer in full serum media **(B)** and HD media **(D).** Images show the initial wound (pink) and migratory cells (orange) in wildtype and D538G mutant cells at 0 hours and 24 hours in full serum media **(C)** and HD media **(E)**. Proliferation and migration figures represent at least 3 independent experiments, performed in triplicate. Error bars represent SEM.

### D538G mutation alters ESR1 genomic binding

To ascertain ESR1 genomic binding sites in Ishikawa cells and the manner in which the D538G mutation alters ESR1’s genomic interactions, we performed ChIP-seq with an antibody that recognizes the FLAG-epitope tag in the two wildtype and three mutant clones after 1 hour treatments with DMSO or E2. We identified less than 55 peaks in each of the two wildtype lines treated with DMSO, indicating negligible ESR1 binding in the absence of E2 treatment. We observed a significant increase in genomic binding, with 20,104 ESR1 binding sites (ERBS) on average identified in the two wildtype cell lines following E2 induction. An analysis of the overlap of binding sites called in the wildtype clones treated with E2 showed 90% (15,780 out of 17,466 on average across replicates) concordance between lines, highlighting the reproducibility of our experimental findings (Figure S3A). To confirm that the FLAG-tag did not significantly alter ESR1 genomic binding, we overlapped the binding sites discovered with the FLAG antibody to binding sites from a ChIP-seq previously done in Ishikawa cells with an antibody that recognizes ESR1 (Gertz 2012; Gertz et al. 2013). We observed 77% overlap (5,583 out of 7,294) on average with this previously generated ESR1 ChIP-seq dataset, indicating that the epitope tag does not drastically affect ESR1 binding in the wildtype cell lines (Figure S3B - E). In contrast, we observed D538G mutant ESR1 binding at more than 22,292 binding sites on average in the three mutant lines in the absence of E2, suggesting that the mutation confers ligand-independent binding in endometrial cancer cell lines.

Analysis of differential binding between wildtype clones treated with E2 and all D538G mutant clone samples established 6,534 constant ESR1 binding sites, 2,205 binding sites that were enriched in the D538G mutant lines, and 3,316 sites that were enriched in the wildtype lines (Figure 4A). When comparing loci enriched in D538G ESR1 binding to loci enriched in wildtype ESR1 binding, we found a 1.30-fold enrichment in intronic regions (p-value = 3.2×10^-6^, Fisher’s exact test), a 1.96-fold depletion in promoter regions (p-value = 1.3×10^-8^, Fisher’s exact test), and a 1.30-fold depletion in intergenic regions (p-value = 4.2×10^-6^, Fisher’s exact test) (Figure S3F). Using the loci enriched in D538G ESR1 binding, *de novo* motif analysis revealed a significant enrichment for the canonical ERE (p-value = 8.5×10^-18^, MEME(Bailey 2009)), indicating that many of these novel sites are direct targets of the D538G mutant (Figure S3G). Additional motifs were also enriched at these sites including the Zic family of transcription factors (p-value = 5.7×10^-16^, MEME), the NF-I transcription factors (p-value = 2.5×10^-17^, MEME) (Bailey 2009) (Bailey 2009), and ebox binding proteins (p-value = 1.7×10^-11^, MEME) (Figure S3G). The sites enriched in ESR1 binding in the wildtype lines were also enriched in the canonical ERE motif (p-value = 1.9×10^-47^, MEME), although other motifs were enriched, including the forkhead factor motif (p-value = 4.1×10^-35^, MEME) and the Sox factor motif (p-value = 6.3×10^-27^, MEME) (Figure S3H). While both mutant-enriched and wildtype-enriched ESR1 bound loci had enrichment of EREs, we found that the strength of EREs found in constant ESR1 binding sites and wildtype-enriched ESR1 binding sites was significantly higher than EREs found in D538G enriched sites (p-value < 2.2×10^-16^, Wilcoxon signed-rank test; Figure 4B). These results indicate that the D538G mutation alters ESR1 genomic binding, including a move to sites with suboptimal EREs.

**Figure 4.**
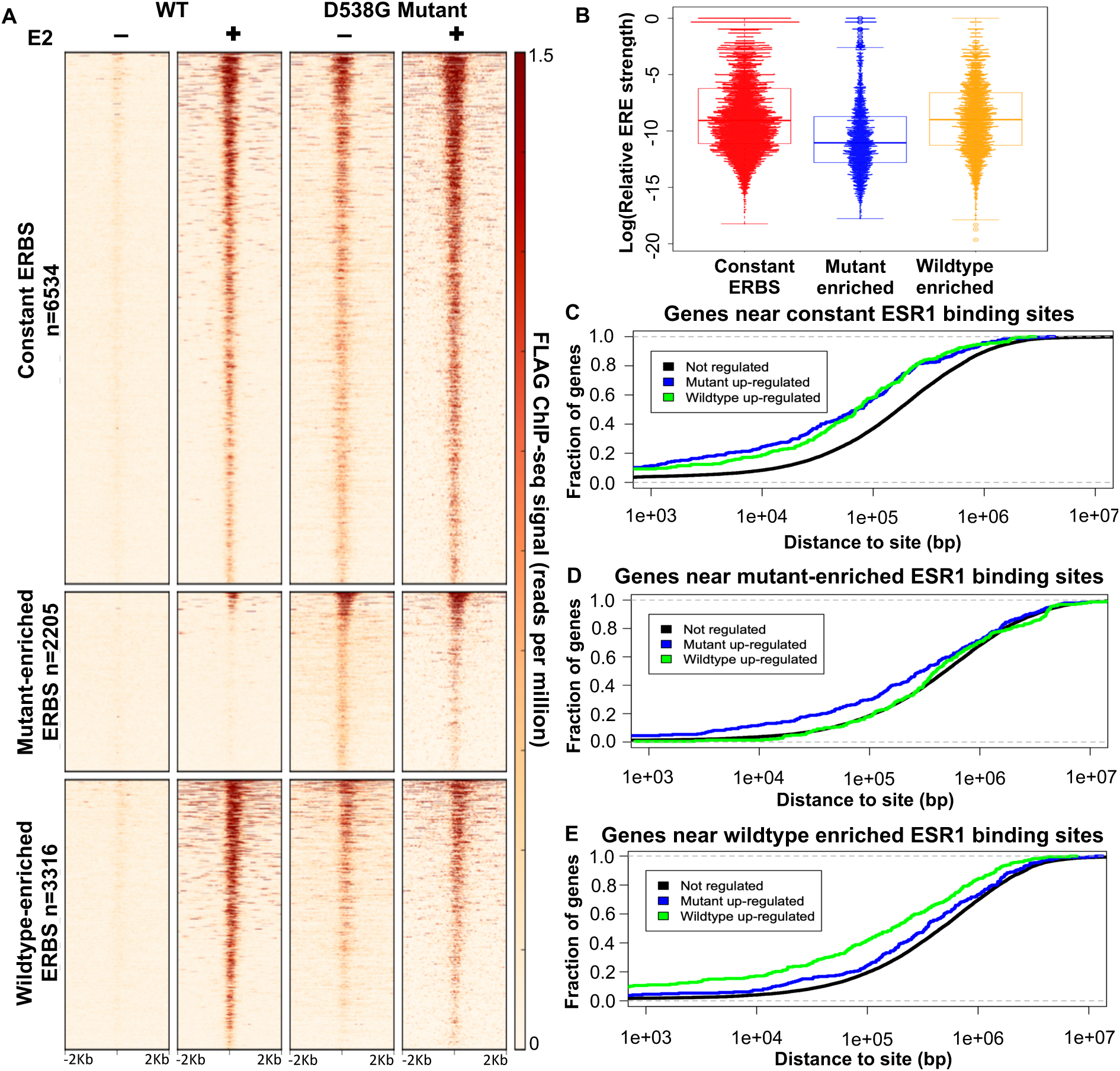
D538G mutation alters ESR1 genomic binding. **A.** Heatmap displays ER binding in representative wildtype and D538G mutant clones, where each row is an ESR1 binding site. The heatmaps include sites that are constant in wildtype and mutant lines (top panels), sites that are enriched in the mutant lines (middle panels) and sites that are enriched in wildtype lines (bottom panels). **B.** Plot shows that the distribution of the predicted relative affinity for ESR1, based on the best match to an ERE, is higher in constant binding sites (red) and wildtype-enriched sites (yellow) than D538G mutant-enriched ESR1 binding sites (blue). Cumulative distribution plots show the fraction of mutant upregulated, downregulated, or not regulated genes that have a constant **(C)**, mutant-enriched **(D)**, or wildtype-enriched **(E)** ESR1 binding site within a given distance from the transcription start site.

The regulatory proteins utilized by ESR1 to mediate gene expression in breast cancer cells have been well characterized by multiple labs (Carroll 2005; Hurtado 2010; Magnani 2011; Tan 2011; Mohammed 2013). However, the proteins responsible for mediating these interactions in endometrial cancer have not been established, but ETV4 (Gertz et al. 2013) and FOXA1 (Droog 2017) have been proposed as key transcription factors. To understand the relationship between mutant ESR1 binding and these factors, we overlapped ESR1 binding sites with the binding sites of FOXA1, which is reported to overlap with 8% of ESR1’s binding sites in parental Ishikawa cells, and ETV4, which is reported to overlap with 45% of ESR1’s binding sites in parental Ishikawa cells (Gertz et al. 2013). We found that mutant-enriched ESR1 binding sites are depleted in FOXA1 binding sites (p-value = 1.0×10^-4^, Fisher’s exact test) compared to wildtype-enriched ESR1 bound sites. In addition, mutant-enriched binding sites were also significantly depleted in ETV4’s binding sites (p-value < 2.2×10^-16^, Fisher’s exact test) compared to wildtype-enriched ESR1 bound sites, suggesting that other transcription factors might be playing a role in mutant ESR1 genomic binding.

To explore the connection between mutant-enriched ESR1 binding and gene expression, we analyzed the distance between mutant-enriched, constant, and wildtype-enriched ESR1 binding sites and the transcription start sites (TSS) of genes upregulated, downregulated or not regulated by the mutation. We found that constant binding sites were closer to both mutant upregulated (p-value = 1.51×10^-15^, Wilcoxon signed-rank test) and mutant downregulated genes (p-value = 6.2×10^-14^, Wilcoxon signed-rank test), than not regulated genes (Figure 4D). Mutant-enriched binding sites were significantly closer to mutant upregulated genes (Figure 4C, p-value = 7.7×10^-15^, Wilcoxon signed-rank test) and wildtype-enriched sites were found closer to genes that were downregulated by the mutation (Figure 4E, p-value = 6.2×10^-16^, Wilcoxon signed-rank test). Overall, the association between genes and ESR1 binding sites suggests that wildtype and mutant ESR1 are generally acting as activators and may be directly contributing to many of the mutant-specific gene expression changes. The association of constant ESR1 binding sites with both mutant upregulated and downregulated genes implies that these ESR1 bound sites may be becoming more or less active when bound by mutant ESR1 compared to wildtype ESR1.

### D538G *ESR1* mutation leads to changes in accessible chromatin

The mutant-specific gene expression changes and ESR1 binding alterations led us to hypothesize that the D538G mutation may affect chromatin accessibility. To test this hypothesis, we performed the Assay for Transposase Accessible Chromatin followed by sequencing (ATAC-seq) in the wildtype and D538G mutant cell lines, after a 1 hour treatment with DMSO or E2. Similar to the distinct RNA-seq profiles, principal component analysis of the isogenic lines once again clustered wildtype and mutant lines separately (Figure 5A). The first principal component accounted for 60% of the variance in the ATAC-seq data and separated wildtype and D538G mutant lines. The second principal component, which accounted for 18% of the variance, separated clones, while E2 treatment for 1 hour was not a large contributor to the variance. This analysis suggests that the 1 hour E2 induction does not have major genome-wide effects on chromatin accessibility in comparison to the D538G *ESR1* mutation.

**Figure 5.**
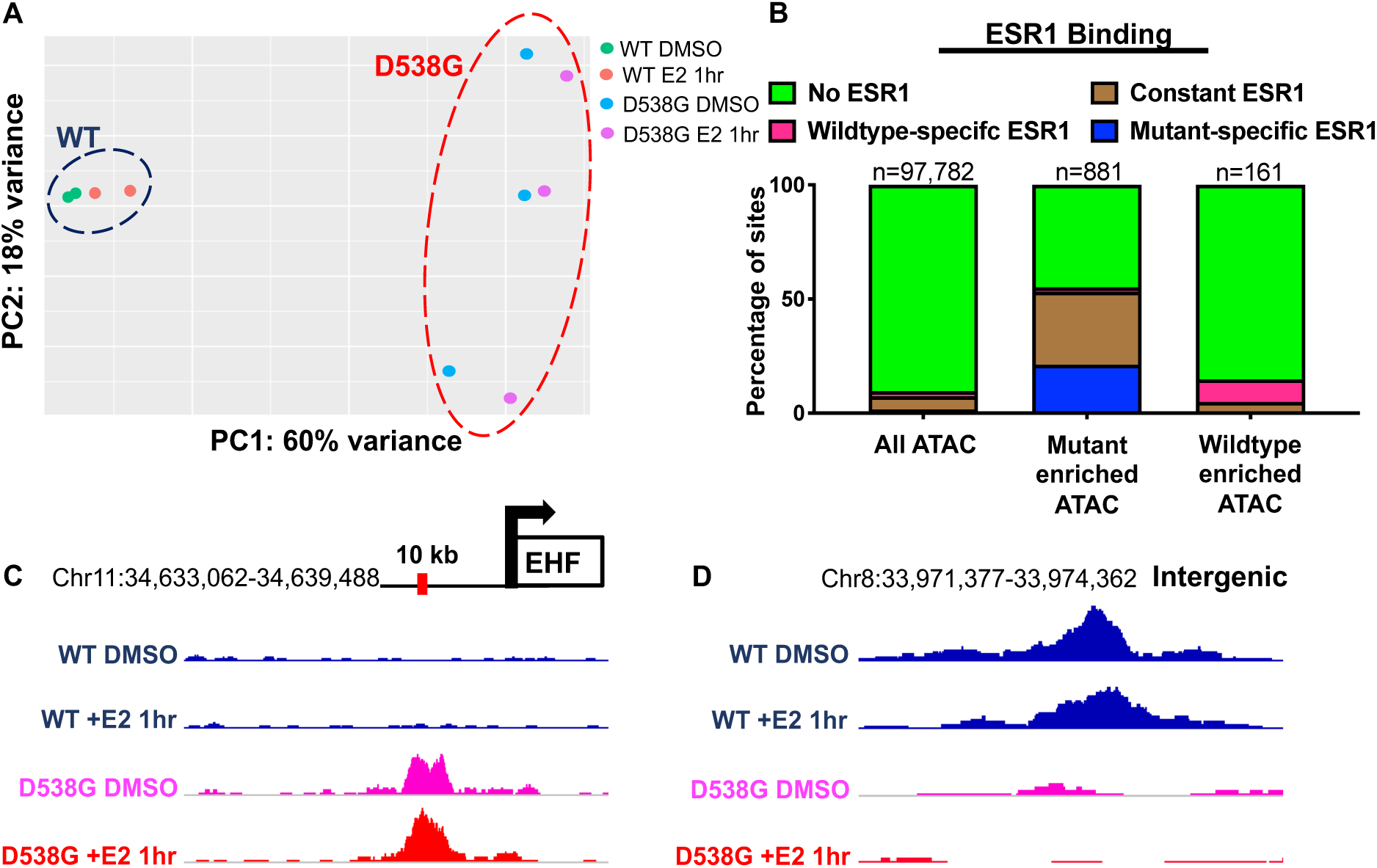
ESR1 D538G mutation alters chromatin accessibility at multiple loci. **A.** Principal component analysis shows the relationship between ATAC-seq signal of ESR1 wildtype (blue circle) and D538G mutant cell lines (red circle). **B.** Less than 10% of All ATAC-seq sites overlap with ESR1 binding sites, while 55% of mutant-enriched ATAC-seq sites overlap ESR1 binding sites, including constant ESR1 binding (brown) and mutant-enriched ESR1 binding (blue). 15% of wildtype-enriched ATAC-seq sites overlap ESR1 binding sites. Representative browser tracks show ATAC-seq signal increases with the D538G mutation at a region near EHF **(C)** and ATAC-seq signal decreases at an intergenic region on Chromosome 8 **(D).** Wildtype ATAC-seq signal DMSO/+E2 (blue), D538G DMSO (pink) and D538G +E2 (red) are scaled to the same value at each locus.

Analysis of ATAC-seq signal from our experiments identified 881 regions that were significantly more accessible and 161 regions that were significantly less accessible in the D538G mutant cell lines compared to wildtype cell lines. Wildtype enriched regions were 14.3-fold more likely to resided in promoter regions compared to mutant enriched regions (p-value < 2.2×10^-16^, Fisher’s exact test), while mutant enriched regions were 1.6-fold more likely to reside in intronic regions (p-value = 0.0098, Fisher’s exact test) (Figure S7B). Examples of these mutant-enriched and mutant-depleted ATAC-seq regions are shown in Figures 5C and 5D and a heatmap of the mutant-enriched sites shows large magnitude changes in ATAC-seq signal at the vast majority of these loci (Figure S5A). We found that while 9.6% of all identified open chromatin regions are associated with ESR1 binding, ESR1 binding was found at the majority of mutant-enriched ATAC-seq sites, with 55% of the mutant-enriched ATAC-seq sites exhibiting ESR1 binding, including constant and mutant-enriched ESR1 binding sites (Figure S5B). Motif analysis at ESR1 associated, mutant-enriched ATAC-seq sites identified the ERE motif as expected (p-value = 3.38×10^-23^, MEME) and the NF-I motif (p-value = 6.64×10^-27^, MEME). These results suggest that mutant ESR1 could be playing a major role in chromatin accessibility, possibly owing to mutant ESR1’s constitutive activity. We found that 45% of mutant-enriched ATAC-seq sites were not associated with ESR1 binding and these sites were enriched for TEAD4 (p-value = 6.41×10^-10^, MEME) and AP-1 (p-value = 4.28×10^-9^, MEME) motifs (Figure S5C), suggesting that other transcription factors may be contributing to mutant-specific alterations in chromatin accessibility.

### Prolonged exposure to estrogen does not recreate mutant-specific regulatory effects

The mechanism by which mutant ESR1 regulates a novel set of genes may be explained by constitutive ESR1 activity or neomorphic functions conferred by the D538G mutation. To determine how much of the gene regulatory effects of mutant ESR1 are due to constitutive ESR1 activity, we cultured the two wildtype clones in the presence of 10 nM E2 for a 25 day period and performed RNA-seq and ATAC-seq on cells collected at 10, 15, 20 and 25 day intervals during this prolonged exposure. Differential gene expression analysis identified 658 genes that were upregulated and 1,138 genes that were downregulated in prolonged E2 treated wildtype cells compared to wildtype cells treated for 8 hours with E2 or DMSO (Figure 6A). These numbers suggest there are more genes regulated by prolonged exposure to E2 than the transient 8 hour induction; however, the overlap with mutant-specific gene expression changes was minimal. Only 25 genes (8.3% of mutant-specific upregulated genes) that were upregulated in response to prolonged E2 were also upregulated by the mutation (Figure 6B, examples in Figure S6). Additionally, only 34 downregulated genes (14% of mutant-specific downregulated genes) overlapped with mutant-specific downregulated genes (Figure 6C). These findings suggest that the D538G mutant regulates a ligand-independent transcriptional program that is dissimilar to prolonged E2 exposure in endometrial cancer cells.

**Figure 6.**
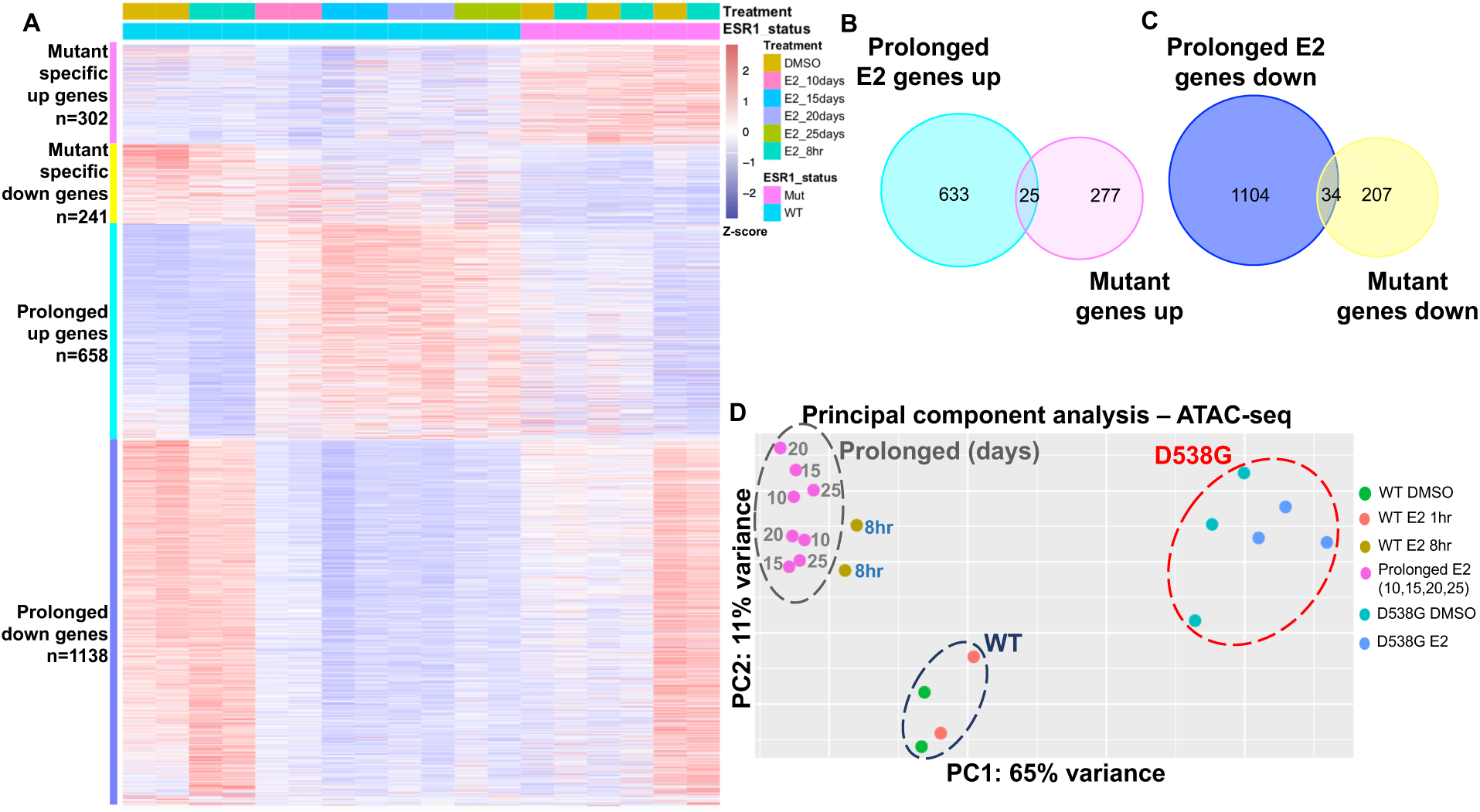
Prolonged E2 exposure does not recapitulate D538G mutant regulatory consequences. **A.** Heatmap shows the relative expression of mutant-specific differentially expressed genes as well as genes up- and down-regulated in response to prolonged E2 (each row is a gene). Samples are indicated by the column annotations described in the legend. **B.** Venn diagram shows the overlap between genes upregulated in wildtype lines exposed to prolonged E2 and mutant-specific upregulated genes. **C.** Venn diagram shows the overlap between genes downregulated in wildtype lines exposed to prolonged E2 and mutant-specific downregulated genes. **D.** Principal component analysis of ATAC-seq signal exhibits four sample groups: wildtype lines with 1 hour or no E2 treatment (navy circle), wildtype lines with 8 hour inductions (blue circle), wildtype lines with prolonged E2 exposure (gray circle, numbers indicate days of treatment), and D538G mutant lines (red circle).

To determine if constitutive ESR1 activity could re-create the mutant-specific chromatin accessibility patterns, we compared the ATAC-seq results from the prolonged E2 experiment in wildtype cells to 1 hour and 8 hour E2 inductions in wildtype cells, as well as the results from the D538G mutant clones. Principal component analysis of all ATAC-seq samples identified three distinct clusters: 1) DMSO controls along with short-term E2 treated wildtype cells (1 hour), 2) prolonged E2 treated wildtype cells (10 days, 15 days, 20 days, and 25 days) along with the 8 hour E2 induction, and 3) the D538G mutant lines (Figure 6D). While Figure 5A indicates that the 1 hour E2 induction did not significantly affect chromatin accessibility globally, prolonged E2 exposure is able to affect chromatin accessibility, with some features being shared with the mutation. However, since the prolonged E2 treated cells do not cluster with the mutant lines, we can conclude that these features do not recapitulate most of the mutant-specific genome-wide effects on chromatin accessibility.

The ATAC-seq analysis revealed 1,488 regions that became more accessible and 2,915 regions that were less accessible following prolonged E2 exposure in wildtype cells (Figure S7). Regions more accessible upon prolonged E2 treatment were 20.8-fold more likely to resided in promoter regions compared to regions less accessible after prolonged E2 treatment (p-value < 2.2×10^-16^, Fisher’s exact test), while less accessible regions were 3.7-fold more likely to reside in intergenic regions (p-value < 2.2×10^-16^, Fisher’s exact test) and 2.5-fold more likely to reside in intronic regions (p-value < 2.2×10^-16^, Fisher’s exact test) (Figure S7C). We overlapped regions that are more accessible after prolonged E2 exposure and regions that are more accessible in the D538G mutant clones, as described above, and identified only 77 regions that were common between datasets, which represented 8.7% of the mutant-enriched regions. Similarly, there were 44 loci that overlapped between chromatin that becomes less accessible in response to prolonged E2 treatment and mutant-depleted accessible chromatin, representing 27% of the mutant-depleted regions (examples in Figure S6). Our findings indicate that prolonged E2 is unable to recapitulate the observed *ESR1* mutant driven changes to chromatin accessibility. Collectively, the effects on gene expression and chromatin accessibility both suggest that the D538G mutation confers distinct neomorphic properties to the receptor which cannot be adequately explained by constitutive estrogen signaling.

## Discussion

Through the creation of isogenic models of a common *ESR1* mutation, D538G, we have found that mutant ESR1 exhibits estrogen-independent activity in endometrial cancer cells. Similar to findings in breast cancer (Merenbakh-Lamin et al. 2013; Robinson et al. 2013; Toy et al. 2013; Jeselsohn et al. 2014; Bahreini 2017; Jeselsohn 2018), mutant ESR1 is able to bind the genome and drive the transcription of estrogen-responsive genes in the absence of estrogens. The estrogen-independent activity of mutant ESR1 that we observed is consistent with the clinical observation that endometrial cancer patients with *ESR1* mutations have lower body mass index than patients without *ESR1* mutations (Backes et al. 2016). Adipose tissue, which is more prevalent in obese patients, is capable of peripheral estrogen production (Siiteri 1987), but endometrial tumors with *ESR1* mutations appear not to rely on this excess estrogen, presumably due to constant ESR1 activity (Rodriguez 2019).

The use of multiple *ESR1* mutant and wildtype clones enabled the discovery of molecular changes that can be reproducibly attributed to the mutation. Wildtype ESR1 binds to different loci in breast cancer and endometrial cancer cells (Gertz et al. 2013) and primary tumors (Droog 2017) leading to different transcriptional responses to E2. Mutant ESR1 exhibits a similar cell type-specific pattern in which the genes regulated by mutant *ESR1* are different between endometrial cancer cells and breast cancer cells. This is true for both estrogen independent regulation of normally estrogen responsive genes as well as novel regulation by mutant ESR1 of non-estrogen responsive genes. These results suggest that mutant ESR1 binding site and target gene selection is still constrained by the other transcription factors and co-factors expressed in the cell as well as the chromatin landscape. Although different genes are regulated by mutant *ESR1* in breast and endometrial cancer cells, similar pathways including cellular growth, proliferation, and movement, are affected by the mutations suggesting that *ESR1* mutations might cause similar phenotypes in breast and endometrial tumors. Motivated by the RNA-seq results, we measured proliferation and found that growth rates were not significantly different between mutant and wildtype clones; however, migration was significantly enhanced in the mutant clones. Consistent with the migration observations, we found that higher expression of several mutant-specific upregulated genes and lower expression of several mutant-specific downregulated genes in our datasets correlate with more aggressive tumors and poorer outcomes for endometrial cancer patients. Together our results indicate that *ESR1* mutations have the potential to drive more dangerous forms of endometrial cancer, which is corroborated by a trend towards worse outcomes for patients with *ESR1* mutant disease (Backes et al. 2016).

*ESR1* mutations do not just confer ligand-independent estrogen signaling. In fact, most of the genes differentially expressed between the mutant lines and the wildtype lines are genes that do not normally respond to E2. This mutant-specific gene regulation appears to be directed by mutant ESR1 with the D538G mutation causing a large alteration in the genomic loci that ESR1 binds. Chromatin accessibility also increases at specific sites across the genome due to the D538G mutation and a majority of these sites are bound by mutant ESR1. The increased chromatin accessibility at some mutant ESR1 bound sites could be the result of different underlying effects. One possibility is a small pioneering role for mutant ESR1, which could be due to changes in co-factor recruitment, as found in breast cancer (Jeselsohn 2018), or because of its constant activity and binding. Another possibility is that ESR1 is taking advantage of the increased activity or expression of another transcription factor (e.g. NF-I factors) that has led to increased chromatin accessibility, which would represent an indirect effect of mutant *ESR1* on the chromatin landscape.

In order to determine how much of mutant *ESR1’*s ability to regulate a new set of genes is related to its constant activity, we treated wildtype clones with continuous saturating doses of E2 for 25 days. Prolonged exposure to E2 changed the expression of thousands of genes and altered chromatin accessibility at thousands of loci; however, there was little overlap with the gene regulatory changes caused by the D538G mutation. These results suggest that the D538G *ESR1* mutation is neomorphic/gain-of-function and does not simply cause hyperactivity. It is unclear how the mutation changes ESR1’s gene regulatory role, but alterations to the placement of helix 12 of the ligand binding domain (Merenbakh-Lamin et al. 2013; Fanning 2016) could cause changes in binding affinities to transcription factors or cofactors that bind to this region. Determining how mutant ESR1 causes novel gene regulation could provide valuable insights into treatment strategies aimed at blocking mutant ESR1’s activity.

In this study, we focused on the D538G mutation because it is the only specific alteration in the LBD; L536 and Y537 have been found to be mutated to several different amino acids (Gaillard 2019). In future studies, it will be interesting to determine if mutations to L536 and Y537 cause similar regulatory and phenotypic changes in endometrial cancer cells, as Y357S and D538G appear to cause mutation specific alterations in breast cancer cells (Bahreini 2017; Jeselsohn 2018). In addition, our study focused on a particular endometrial cancer cell line, Ishikawa, and the effects of *ESR1* mutations may be different in different models. We have recently found that ESR1 genomic binding is consistent between endometrial tumors and distinct from breast tumors, with Ishikawa exhibiting a clear endometrial cancer ESR1 binding pattern (Rodriguez et al. 2019), suggesting that our findings could be generally applicable. In summary, our study has led to the creation of isogenic models of mutant *ESR1* in endometrial cancer cells, the confirmation of estrogen-independent mutant ESR1 activity, and the discovery of novel gene regulation through mutant ESR1 that cannot be explained by constant activity alone.

## Materials and Methods

### Plasmid construction for *ESR1* LBD mutant generation

Mutant cell lines were created using the CETCH-seq method (Savic et al. 2015) in which a pFETCH plasmid is the homology donor (containing the mutation, 3x FLAG tag, P2A linker, and Neomycin resistance gene) and Cas9 is targeted proximal to the stop codon by guide RNAs. To create the pFETCH homology donor plasmid with the D538G *ESR1* LBD mutation, we used primers (see Table S3 for sequences) to PCR amplify *ESR1* homology arms, using a gBlock (IDT, Table S3) that encompassed 1000 bp surrounding the D538G mutation as a template for the amplification of homology arm 1 and genomic DNA from Ishikawa cells (Sigma-Aldrich) as the template for homology arm 2. For amplification, Phusion high fidelity master mix (New England BioLabs) was used with 10 μM of each primer and 1 ng gBlock or 50 ng genomic DNA and amplified for 25 cycles. To generate the wildtype pFETCH donor plasmid, we repeated this technique with Ishikawa genomic DNA as template for arm 1 amplification. Using BsaI and BbsI (New England BioLabs), we double digested the destination pFETCH vector (Addgene 63934, a gift from Eric Mendenhall & Richard M. Myers) and used Gibson assembly HiFi master mix (New England BioLabs) to clone homology arms 1 and 2 simultaneously to create D538G or wildtype pFETCH plasmids. Clones for each pFETCH vector underwent minipreps (Zymo Research) and were verified by Sanger sequencing (Genewiz). In order to create a Cas9 and guide RNA expressing plasmid that targeted near the stop codon of *ESR1*, a Cas9 and guide RNA expression vector (Addgene 62988, a gift from Feng Zhang) was digested with BbsI (New England BioLabs). Guide RNA oligos (Table S3) were annealed and then ligated into the Cas9 guide RNA vector. Clones for each guide RNA underwent minipreps (Zymo Research) and were verified by Sanger sequencing (Genewiz). The pFETCH mutant and wildtype plasmids were SalI (New England BioLabs) digested prior to transfection to linearize the vector.

### Cell culture and transfection for generation of *ESR1* LBD mutant and wildtype lines

Ishikawa cells (Sigma-Aldrich) were seeded in 6-well plates at a density of 300,000 cells per well in RPMI-1640 (Thermo Fisher Scientific) with 10% fetal bovine serum (Thermo Fisher Scientific) and 1% penicillin-streptomycin (Thermo Fisher Scientific). At approximately 50% confluency, 250 ng of each of the two Cas9 guide RNA vectors and 250 ng of either the mutant or wildtype pFETCh vectors were transfected into cells with Lipofectamine 3000 (Thermo Fisher Scientific). Cells were also treated with 1 μM SCR7 (Xcessbio) to inhibit non-homologous end joining (NHEJ) for 3 days post-transfection. 72 hours post-transfection, the media was changed and G418 (Thermo Fisher Scientific) was added to a final concentration of 200 μg/mL. RPMI media and G418 were replaced every 2 days until resistant cells remained. To generate single-cell colonies, transfected cells were plated at limiting dilution and cultured until colonies were large enough to identify visually. Individual colonies were picked and transferred to a 24-well plate and grown until they reached confluency. At this time, genomic DNA was extracted from individual colonies using the DNeasy Blood and Tissue Kit (Qiagen). To genotype clones, we amplified a 700 bp region within the ligand binding domain of *ESR1* using ESR1-LBD_F4 and ESR1-LBD_R1 primers (Table S4). PCR products were purified with Ampure XP beads (Beckman Coulter) and Sanger sequenced (Genewiz) to confirm the D538G mutation or wildtype sequence (Table S5). We also verified the presence of a tagged and untagged copy of *ESR1* using ESR1-LBD_F4 and ESR1-LBD_STOP_R1 primers (Table S4). The wildtype and *ESR1* LBD mutant lines all harbored both a tagged and untagged copy of wildtype *ESR1*. Positive clones then underwent immunoblotting (see below) to verify protein expression.

### Immunoblotting

Ishikawa *ESR1* LBD mutant and wildtype cell lines were seeded in 100mm dishes with RPMI-1640 with 10% fetal bovine serum and 1% penicillin-streptomycin until they reached confluency. At confluency, cells were washed with cold phosphate buffered saline (PBS), adherent cells were scraped, lysed with RIPA buffer (1X PBS, 1% NP-40, 0.5% sodium deoxycholate, 0.1% SDS) supplemented with protease inhibitors (Thermo Fisher Scientific) and sonicated with an Active Motif EpiShear probe-in sonicator with 3 cycles of 10 seconds on, 10 seconds of rest at 40% amplitude. 80 μg of protein was loaded onto a 4-12% Bis-Tris gel (Thermo Fisher Scientific) and electrophoresed for 90 minutes at 130V in 1X MOPS buffer (Thermo Fisher Scientific). Proteins were transferred onto a polyvinylidene difluoride (PVDF) membrane with the iBlot system (Thermo Fisher Scientific). Membranes were blocked in 5% milk/PBST for 1 hour at room temperature. Blots were probed with primary antibodies to FLAG 1:1,000 (Sigma-Aldrich M2), ESR1 1:200 (Santa Cruz HC-20) and actin beta 1:1,000 (Santa Cruz C4) in 2.5% milk/PBST overnight at 4° C. Membranes were washed with PBST and incubated in secondary antibody (Goat anti-Mouse IgG, Goat anti-Rabbit IgG; Thermo Fisher Scientific) in PBST at 1:5,000 dilution. Protein signal was detected with SuperSignal West Femto Maximum Sensitivity Substrate (Thermo Fisher Scientific).

### Cell culture

Ishikawa *ESR1* LBD mutant and wildtype cells were cultured in full media: RPMI-1640 with 10% fetal bovine serum and 1% penicillin-streptomycin. Cells were incubated at 37°C with 5% CO2 for the duration of all experiments. At least 5 days prior to inductions, cells were placed in hormone deprived media: phenol-red free RPMI-1640 media (Thermo Fisher Scientific), since phenol-red is estrogenic, with 10% charcoal-dextran stripped fetal bovine serum (Thermo Fisher Scientific) and 1% penicillin-streptomycin. Media was changed 1 day prior to inductions and cells were treated with either DMSO (vehicle) or 10 nM E2 for 1 hour for ChIP-seq and ATAC-seq experiments, or 8 hours for RNA-seq and ATAC-seq experiments. For prolonged estrogen RNA-seq and ATAC-seq experiments, wildtype cell lines were cultured in phenol-red free RPMI with 10% charcoal-dextran stripped fetal bovine serum and 1% pen-strep for up to 25 days. Cells were treated with 10 nM E2 every 2 days with media changes, and cell lysates were collected at the following time points: 8 hours, day 10, day 15, day 20, day 25.

### ERE luciferase reporter assay

Approximately 15,000 cells per well for each cell line were seeded in a 96-well plate in hormone deprived media on day 5. Cells were then transfected, according to the manufacturer’s protocol with inducible dual-luciferase *and Renilla* estrogen response element (ERE) constructs (Qiagen) using FuGENE HD transfection reagent (Promega). One day post-transfection, media was changed and cells were treated with either DMSO or 10 nM E2. 24 hours post-treatment, luciferase activity was measured using the Dual-Glo luciferase assay system (Promega) as per manufacturer’s instructions. All experiments were performed in individual wildtype and D538G mutant clones in triplicate with three biological replicates. Statistical analysis was performed using Student’s *t*-test.

### Quantitative PCR

Cell lysates were harvested following an 8 hour E2 or DMSO control induction with buffer RLT plus (Qiagen) containing 1% beta-mercaptoethanol (Sigma-Aldrich). Total RNA was extracted and purified using a Quick RNA Mini Prep kit (Zymo Research). 25 ng of RNA was used as starting material per sample and qPCR was performed using the Power SYBR Green RNA-to-CT 1-Step Kit (Thermo Fisher Scientific) on a CFX Connect Real-Time light cycler (Bio-Rad). Primers for *PGR, MMP17, EHF, EPHA3*, and *CTCF* are listed in Table S6. Expression measurement were calculated with the ΔΔC_t_ method using *CTCF* as a control. Experiments were performed in all wildtype and D538G mutant clones in triplicate.

### Proliferation and migration assays

Ishikawa *ESR1* LBD mutant and wildtype cells were cultured in full media or hormone deprived media for 3 days prior to plating for proliferation and migration experiments. For proliferation experiments, approximately 5,000 cells per well for each cell line were seeded in 96 well plates in both media conditions. Cell proliferation was monitored via time-lapse image acquisition every 2 hours, for up to 72 hours, via the IncuCyte ZOOM imaging platform (Sartorius). Doubling times for individual cell lines in the two media conditions were calculated by performing linear regression between hours and the log base 2 confluency percentages that were normalized to the starting confluency. Doubling times were taken as 1 divided by the slope of the best fit line. Migration experiments were assessed via the wound healing assay. Approximately 30,000 wildtype and D538G mutant cells were seeded in 96 well ImageLock Microplates (Sartorius) and grown to confluency in full media and hormone deprived media. The cell monolayer was scraped and migration was monitored via time-lapse image acquisition every 2 hours via the IncuCyte ZOOM imaging platform (Sartorius). Migration rates were calculated as relative wound density over time for wildtype and mutant cell lines. All experiments were performed in individual wildtype and D538G mutant clones in triplicate with three biological replicates. Statistical analysis was performed using Student’s *t*-test.

### ChIP-seq

Ishikawa *ESR1* mutant and wildtype clonal cell lines were induced with DMSO or 10 nM E2 for one hour, followed by fixation with 1% formaldehyde for 10 minutes at room temperature to cross-link cells. The cross-linking reaction was stopped with the addition of glycine to a final concentration of 125 mM. Cells were washed with cold PBS and harvested via cell scraping in Farnham lysis buffer supplemented with protease inhibitors. Chromatin immunoprecipitation was performed as previously described (Reddy 2009) with an Anti-FLAG (Sigma-Aldrich M2) antibody. ChIP-seq libraries were sequenced on an Illumina HiSeq 2500 and sequencing reads were aligned to the hg19 build of the human genome using Bowtie (Langmead 2009) with the following parameters: -m 1 -t --best -q -S -I 32 -e 80 -n 2. The hg19 build of the human genome was used for all genomic analyses. We do not believe that realigning reads to the current genome build (GRCh38) would substantially change results, since we are restricting our analyses to uniquely alignable regions of the genome. MACS2 (Zhang 2008) was used to call peaks with a p-value cutoff of 1×10^-10^ and a mfold parameter between 15 and 100. Input control libraries from both wildtype and mutant cell lines were used as controls for each ChIP-seq experiment. All ChIP-seq experiments were done in biological duplicates. Genomic annotation of binding sites was performed using CEAS (Ji et al. 2006). Overlaps between peaks were determined using a 1 basepair minimum overlap and percentages were calculated by dividing the number of overlapping peaks by the number of peaks in the smaller set (i.e. the percentage of maximal possible overlap). Differential binding sites were identified by using DESeq2 (Love 2014) to compare counts per million at each ERBS that was identified in any ChIP-seq sample. ESR1 ChIP-seq experiments in wildtype clones treated with E2 were compared to all ESR1 ChIP-seq experiments performed in mutant clones. An adjusted p-value cutoff of 0.05 was used to identify mutant-enriched and wildtype-enriched ESR1 bound sites. Constant ERBS were regions bound by ESR1 in at least one replicate of the wildtype E2 FLAG ChIP-seq experiments and at least one replicate of the mutant FLAG ChIP-seq experiments, but were not found to be differentially bound in the DESeq2 analysis. Motif finding was performed on 500bp regions surrounding the summit of identified peaks. Motifs between 6 and 30 basepairs in length were identified by MEME Suite (Bailey 2009), with a motif distribution of zero to one occurrence per sequence. To identify strong and weak EREs in mutant-specific and constant ESR1 binding sites, ERE scores were calculated by Patser (Hertz 1999), within 100bp of an individual peak’s summit.

### RNA-seq

Following 8 hour treatments in hormone deprived media with either 10 nM E2 or DMSO, cells were washed with PBS and harvested with buffer RLT plus (Qiagen) containing 1% beta-mercaptoethanol (Sigma-Aldrich). Cells were passed through a 21-gauge needle and syringe (Sigma-Aldrich) to lyse genomic DNA before RNA was extracted and purified using a Quick RNA Mini Prep kit (Zymo Research). Using the KAPA Stranded mRNA-Seq kit (KAPA Biosystems), poly(A) selected libraries were created with 500 ng of input RNA per sample. RNA-seq libraries were sequenced on an Illumina HiSeq 2500 and sequencing reads were aligned to the hg19 build of the genome using HISAT2 (Kim 2015), with The University of California Santa Cruz (UCSC) Known Genes definitions used to build indexes. Sam files were converted to BAM files and sorted with SAMtools (Li 2009). To quantify reads that mapped to UCSC Known Genes, we used featureCounts (Liao 2014). Reads were normalized and analyzed for differential enrichment using the DESeq2 package in R (Love 2014). RNA-seq experiments were done in each wildtype and D538G mutant clone, and clones with the same genotype were used as biological replicates in the analysis. To parse differentially regulated gene lists, we first identified statistically significant genes (adjusted p-value < 0.05) that were upregulated and downregulated in response to an E2 induction in wildtype cells. To find genes that are regulated by E2 in wildtype clones and change expression in the mutant clones without E2, we used DESeq2 to compare wildtype DMSO treated samples to mutant DMSO treated samples and compared the genes to E2 regulated genes in wildtype clones. To identify novel genes not normally regulated by E2, we used DESeq2 to compare all D538G samples to all wildtype samples treated with E2 and then subtracted genes that were regulated by E2 in wildtype cells. The adjusted p-value cutoff for all significant genes in each comparison was < 0.05. Differentially expressed novel upregulated and downregulated genes were analyzed through the use of Ingenuity Pathway Analysis (QIAGEN Inc., https://www.qiagenbioinformatics.com/products/ingenuity-pathway-analysis). The five statistically significant molecular and cellular functions identified in this analysis are included in the supplemental data (Figure S1B,C).

For prolonged RNA-seq experiments *ESR1* wildtype cell lines were treated with 10 nM E2 every 2 days along with media changes and cell lysates were collected at the following time points: day 10, day 15, day 20, and day 25. Total RNA was extracted, poly(A) selected libraries were constructed, sequenced, and analyzed as described above. The two wildtype clones were used as biological replicates in this analysis. To identify genes differentially regulated in response to prolonged E2, we compared a group consisting of the DMSO treated and 8 hour E2 treated samples to a group consisting of the 10, 15, 20, and 25 day samples. For DESeq2 analysis, the two groups were treated as categorical variables. The adjusted p-value cutoff for all significant genes was < 0.05.

### TCGA Data Analysis

RNA-seq and clinical data was obtained from the TCGA data portal in December 2015. The gene expression measurements used were level 3 RNaseqV2 normalized RSEM data. Only samples with endometrioid histology and *ESR1* expression above the median among the endometrioid tumors were analyzed for survival analysis. Cox regression was used to evaluate the association between gene expression and progression-free survival in R using coxph from the survival package. We used a median cutoff to define high and low expressing tumors for each gene when running the Cox regression.

### ATAC-seq

After 1 and 8 hour treatments with either 10 nM E2 or DMSO, cells were trypsinized and isolated by centrifugation. 250,000 cells were isolated from *ESR1* LBD wildtype or mutant cell lines and ATAC-seq was performed as previously described (Buenrostro 2013). ATAC-seq libraries were sequenced on an Illumina HiSeq 2500 and sequencing reads were aligned to hg19 using Bowtie (Langmead 2009) with the following parameters: -m 1 -t --best -q -S -I 32 -e 80 -n 2. Sam files were converted to BAM files and sorted with SAMtools (Li 2009). MACS2 (Zhang 2008) was used to call peaks without an input control but with a p-value cutoff of 1*10^-10^. We used featureCounts (Liao 2014) to quantify reads that aligned to all ATAC-seq peaks called in any sample. Reads were normalized and analyzed for differential enrichment using the DESeq2 package for R (Love 2014). ATAC-seq experiments were done in each wildtype and D538G mutant clone, and clones with the same genotype were used as biological replicates. Mutant-enriched and mutant-depleted ATAC-seq sites were identified by comparing ATAC-seq signal between all wildtype and all mutant samples. Motif discovery at ESR1 associated and non-ESR1 associated regions was performed on 500bp regions surrounding the summit of identified peaks. Motifs between 6 and 30 basepairs in length were identified by MEME Suite (Bailey 2009), with a motif distribution of zero to one occurrence per sequence.

For prolonged ATAC-seq experiments *ESR1* wildtype clones were treated with 10 nM E2 every 2 days during media changes and cells were collected at the following time points: day 10, day 15, day 20, and day 25. ATAC-seq experiments and analysis were performed as described above. Prolonged E2-enriched and depleted ATAC-seq regions were identified by comparing ATAC-seq signal between all wildtype samples treated with DMSO or E2 for 1hr to all wildtype samples exposed to prolonged E2 (10 day, 15 day, 20 day, and 25 day). For DESeq2 analysis, the two groups were treated as categorical variables. The adjusted p-value cutoff for all significant genes was < 0.05. Genomic annotation of these regions was performed using CEAS (Ji et al. 2006).

### Statistical and graphical packages

The statistical analyses were performed in R version 3.3.2 (R Core Team 2017), except for the p-values for novel gene enrichments calculated by IPA and p-values for enriched motifs calculated by MEME. Heatmaps were generated in R using the pheatmap package, and statistical tests used for analysis and corresponding p-values can be found throughout the text. Heatmaps for ChIP-seq and ATAC-seq data were generated by displaying the z-score across a region based on the reads per million that aligned to each region in each sample.

## Data Access

All raw and processed sequencing data generated in this study have been submitted to the NCBI Gene Expression Omnibus (GEO; http://www.ncbi.nlm.nih.gov/geo/) under the following accession numbers: GSE132428 (RNA-seq), GSE132426 (ChIP-seq), and GSE132424 (ATAC-seq).

## Author Contributions

Conceptualization, Z.B. and J.G.; Methodology, Z.B. and J.G.; Investigation, Z.B., K.C.B., S.A. and J.G.; Formal Analysis, Z.B., J.M.V. and J.G.; Writing – Original Draft, Z.B. and J.G.; Writing – Review & Editing, Z.B., J.M.V., K.C.B., S.A., and J.G.; Funding Acquisition, J.G.

## Acknowledgments

This work was supported by a DOD Breast Cancer Research Program Breakthrough Award to J.G., and the Huntsman Cancer Institute. Research reported in this publication utilized the High-Throughput Genomics Shared Resource at the University of Utah and was supported by NIH/NCI award P30 CA042014. We thank Ed Grow for providing reagents, Margit Janat-Amsbury, Jennifer Richer, K-T Varley, as well as Gertz and Varley laboratory members for their helpful comments on the study and the manuscript.

