## Supplemental Figures and Tables S3-6 for "Estrogen-independent molecular actions of mutant estrogen receptor alpha in endometrial cancer"

A

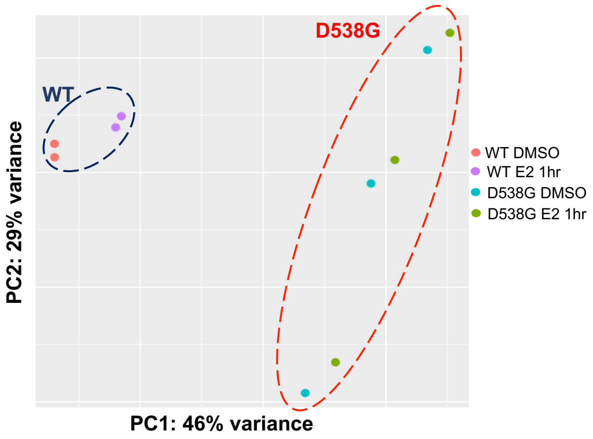


B

| Molecular & Cellular Functions (Up) | p-value |
|---|---|
| Cellular Development | 2.54E-02 – 1.65E-04 |
| Cellular Growth & Proliferation | 2.54E-02 – 1.65E-04 |
| Cellular Movement | 2.54E-02 – 4.84E-04 |
| Cell Cycle | 2.54E-02 – 5.37E-04 |
| Cell-To-Cell Signaling & Interaction | 2.54E-02 – 6.25E-04 |

C

| Molecular & Cellular Functions (Down) | p-value |
|---|---|
| Cellular Movement | 8.45E-03 – 8.06E-08 |
| Cellular Assembly and Organization | 7.50E-03 – 1.76E-06 |
| Cellular Function and Maintenance | 8.66E-03 – 3.26E-05 |
| Cellular Development | 2.96E-03 – 6.46E-05 |
| Cellular Growth and Proliferation | 6.18E-03 – 6.46E-05 |

**Supplemental Figure 1. Gene expression changes due to D538G mutation (related to Figure 2). A.** Principal component analysis of gene expression levels shows the relationship between *ESR1* wildtype (blue circle) and D538G mutant cell lines (red circle) in hormone deprived media treated with DMSO or E2 for eight hours. Ingenuity Pathway Analysis identified molecular and cellular functions enriched in mutant-specific upregulated genes (**B**) and mutant-specific downregulated genes (**C**).

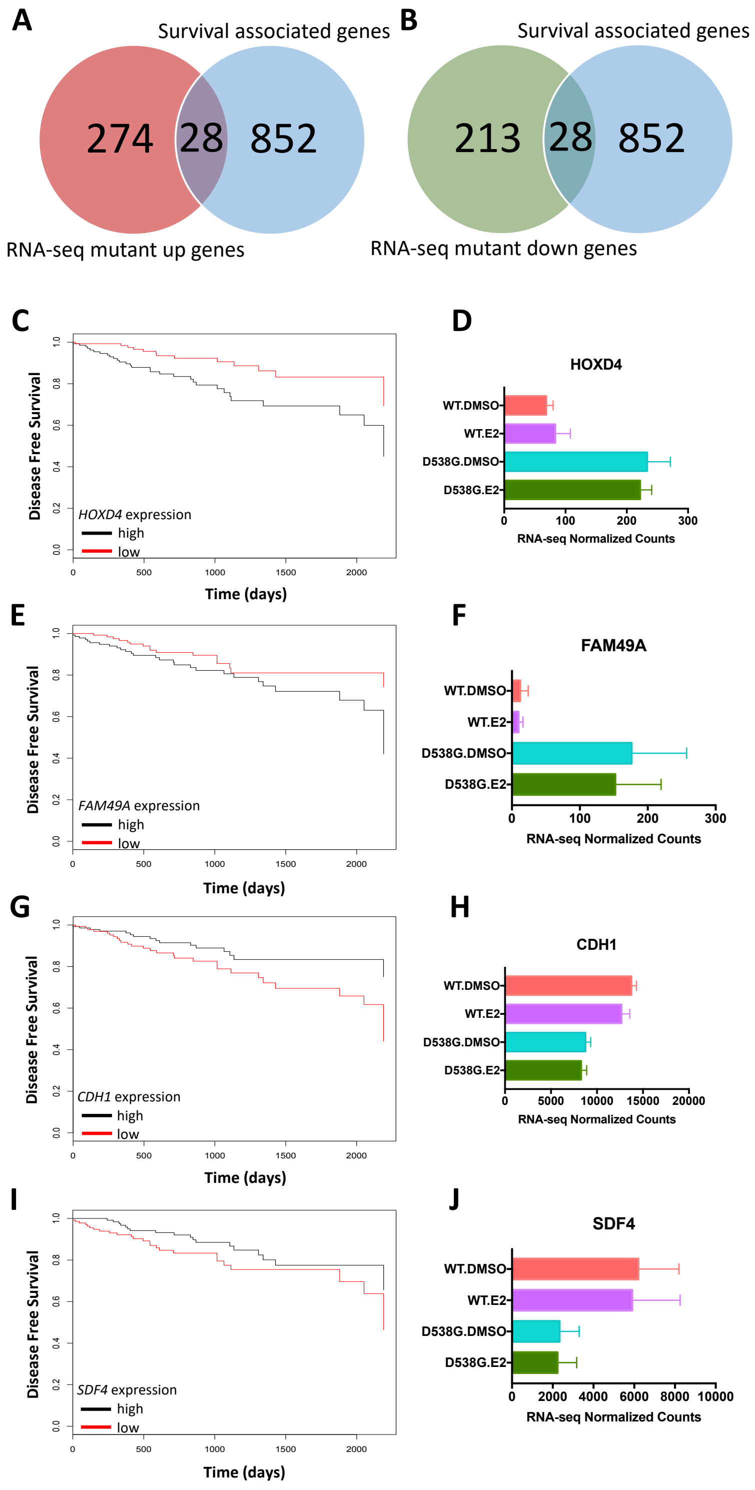


**Supplemental Figure 2. Association between mutant-specific gene expression and endometrial cancer patient outcomes (related to Figure 2).** Venn diagrams show the overlap between genes whose expression is associated with survival in TCGA data and mutant-specific upregulated **(A)** or downregulated genes **(B)**. Examples of Kaplan-Meier plots **(C)** and **(E)** showing the association between disease free survival in endometrial cancer patients and **(D)** *HOXD4* and **(F)** *FAM49A* mutant-specific up-regulated gene expression, where higher expression (black line) is correlated with worse prognosis. Examples of Kaplan-Meier plots **(G)** and **(I)** showing the association between disease free survival in endometrial cancer patients and **(H)** *CDH1* and **(J)** *SDF4* mutant-specific down-regulated gene expression, where higher expression (black line) is correlated with better prognosis.

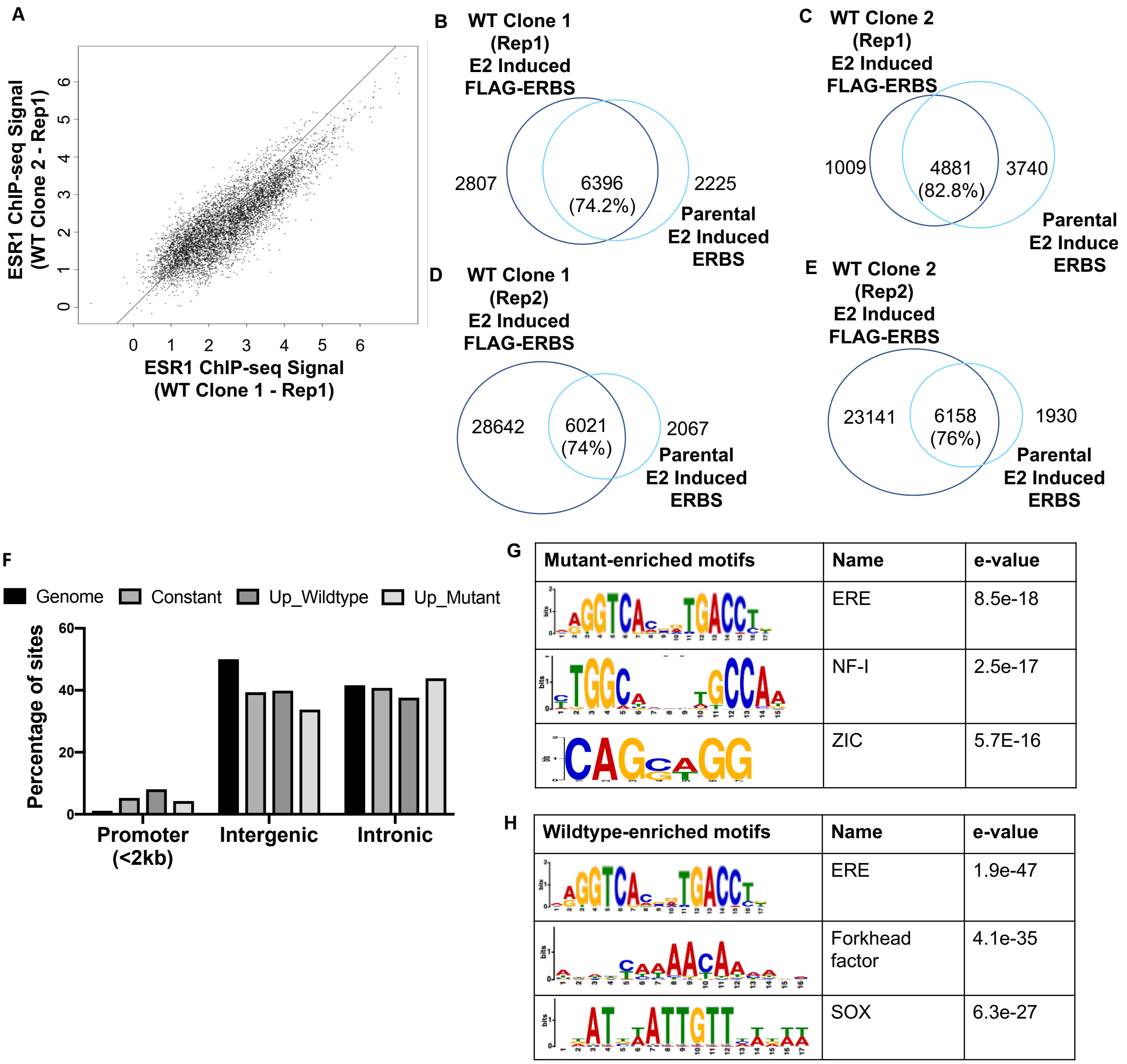


**Supplemental Figure 3. Analysis of FLAG-tagged ESR1 ChIP-seq (related to Figure 4). A.** Scatterplot showing representative overlaps between FLAG-tagged ESR1 binding sites (ERBS) called in two ESR1 wildtype clones (replicate 1) following a one hour E2 induction. **B.** FLAG-tagged ERBS called in wildtype clone one (rep 1) and an ESR1 ChIP done in parental Ishikawa cells following a one hour E2 induction, **(C)** FLAG-tagged ERBS called in wildtype clone two (rep 1) and an ESR1 ChIP done in parental Ishikawa cells following a one hour E2 induction. **D.** FLAG-tagged ERBS called in wildtype clone one (rep 2) and an ESR1 ChIP done in parental Ishikawa cells following a one hour E2 induction, **(E)** FLAG-tagged ERBS called in wildtype clone two (rep 2) and an ESR1 ChIP done in parental Ishikawa cells following a one hour E2 induction. **F.** Genomic distribution of promoters, intergenic and intronic regions in constant, wildtype-enriched and mutant-enriched ESR1 binding sites. Significantly enriched motifs found in mutant-enriched ESR1 binding sites **(G)** and wildtype-enriched ESR1 binding sites **(H)**.

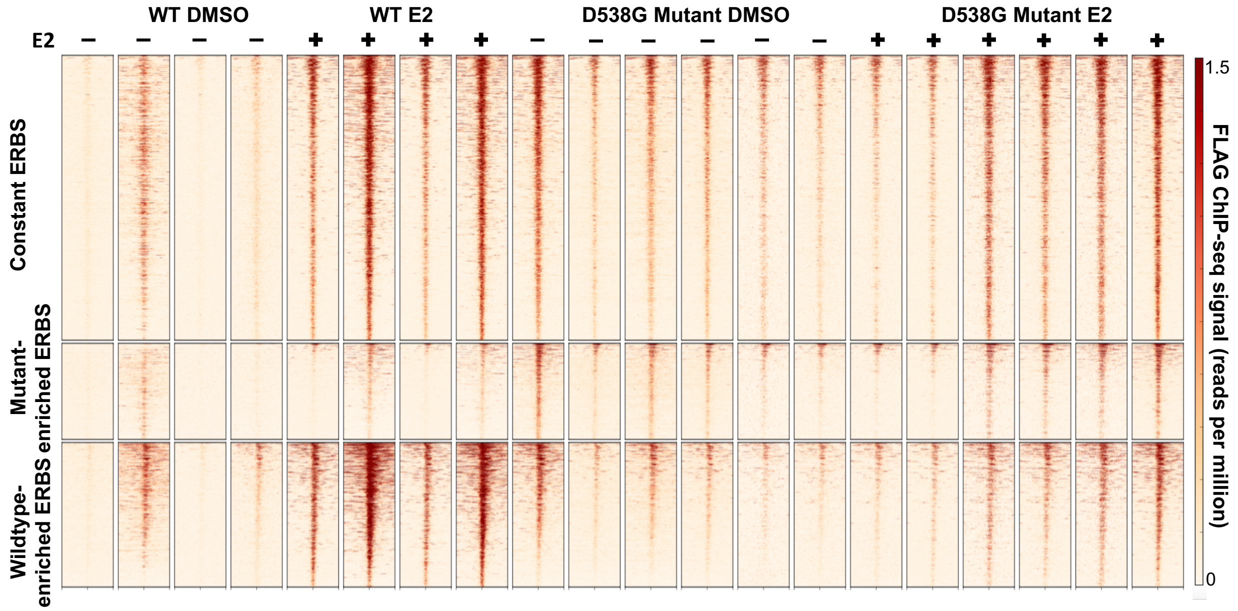


**Supplemental Figure 4. ESR1 bound sites in mutant and wildtype lines (related to Figure 4).** Heatmaps display wildtype and D538G mutant binding in hormone deprived media following treatment with DMSO or E2 for one hour. ESR1 binding sites include constant regions that are similar in wildtype and mutant lines (top panels), sites that are enriched in the mutant lines (middle panels), and sites that are enriched in wildtype lines (bottom panel). Each line is a an ERBS site, regions represent 2.5kb up- and down-stream of the peak summit.

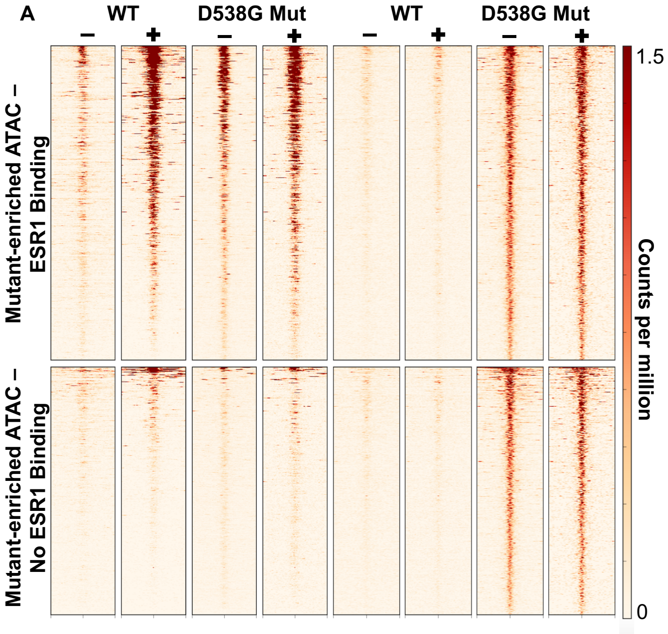


**B**

| Mutant-enriched with ESR1 binding overlap | Name | e-value |
|---|---|---|
| 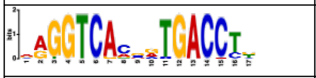 | ERE | 3.38e-23 |
| 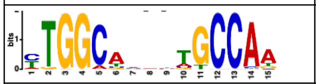 | NF-I | 6.64e-27 |

**C**

| Mutant-enriched with no ESR1 binding overlap | Name | e-value |
|---|---|---|
| 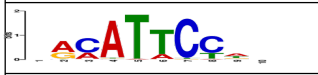 | TEAD4 | 6.41e-10 |
| 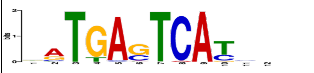 | AP-1 | 4.28e-9 |

**Supplemental Figure 5. Analysis of ATAC-seq regions and ESR1 binding (related to Figure 5). A.** Heatmap displays mutant-enriched ATAC-seq regions that overlap with ESR1 binding (top panels) and regions that do not overlap with ESR1 binding (bottom panels) in representative wildtype and D538G mutant cell lines. Each row is a region that is 2.5kb up- and down-stream of a peak summit. Significantly enriched motifs found in mutant-enriched ATAC-seq sites that overlap with ESR1 **(B)** and motifs enriched in regions that do not overlap with ESR1 **(C)**.

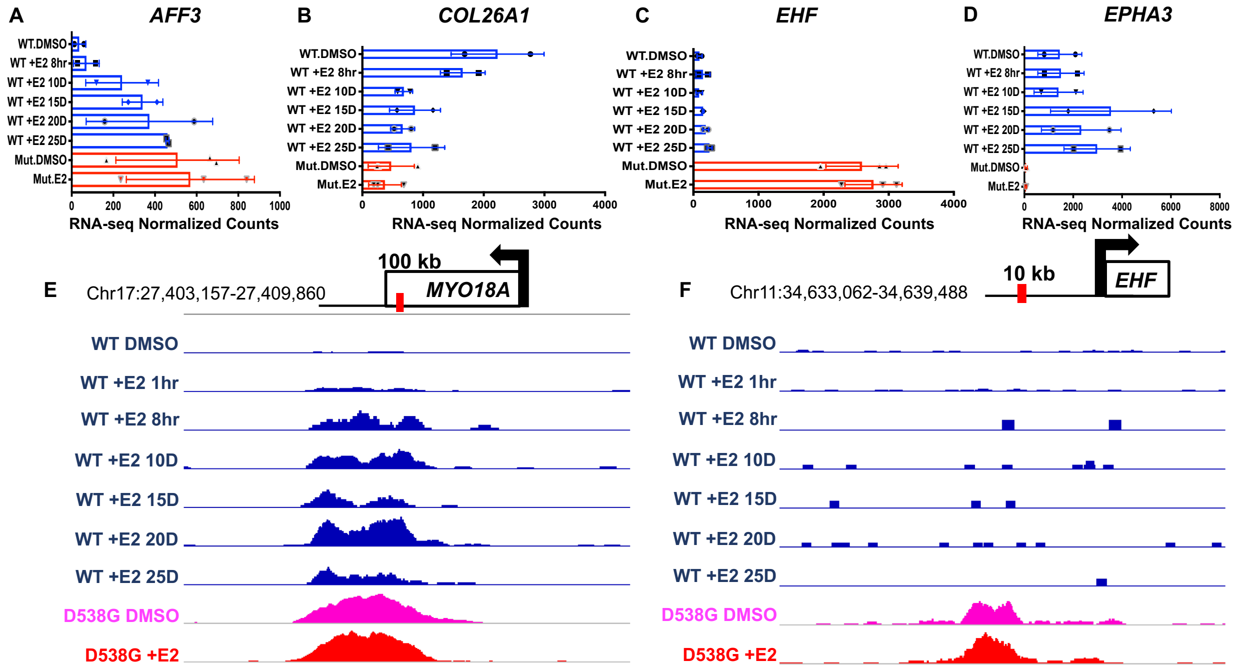


**Supplemental Figure 6. Examples of prolonged E2 exposure effects (related to Figure 6).** Examples of novel mutant-specific regulated genes: **(A)** *AFF3* is upregulated in response to prolonged E2, similar to expression levels seen with the D538G mutation, **(B)** *COL26A1* is downregulated in response to prolonged E2, similar to expression levels seen with the D538G mutation, **(C)** *EHF* expression levels do not change in response to prolonged E2, however this gene is highly expressed in D538G mutant lines, **(D)** *EPHA3* has variable expression levels in response to prolonged E2 but is not expressed in D538G mutant lines. All expression figures show average RNA-seq normalized counts for two wildtype and three ESR1 D538G mutant clones. **E.** Representative browser tracks indicating ATAC-seq signal at loci 100kb downstream of *MYO18A,* a region of more accessible chromatin in response to prolonged E2 exposure. This region is also open in the D538G mutant lines. **F.** Representative browser tracks indicating ATAC-seq signal at loci 10kb upstream of *EHF*, a region of less accessible chromatin in response to prolonged E2 exposure. This region is open in the D538G mutant lines. For the browser tracks, wildtype ATAC-seq signal DMSO/+E2/Prolonged E2 (blue), D538G DMSO (pink) and D538G +E2 (red) are scaled to the same value within each region. Error bars represent SEM.

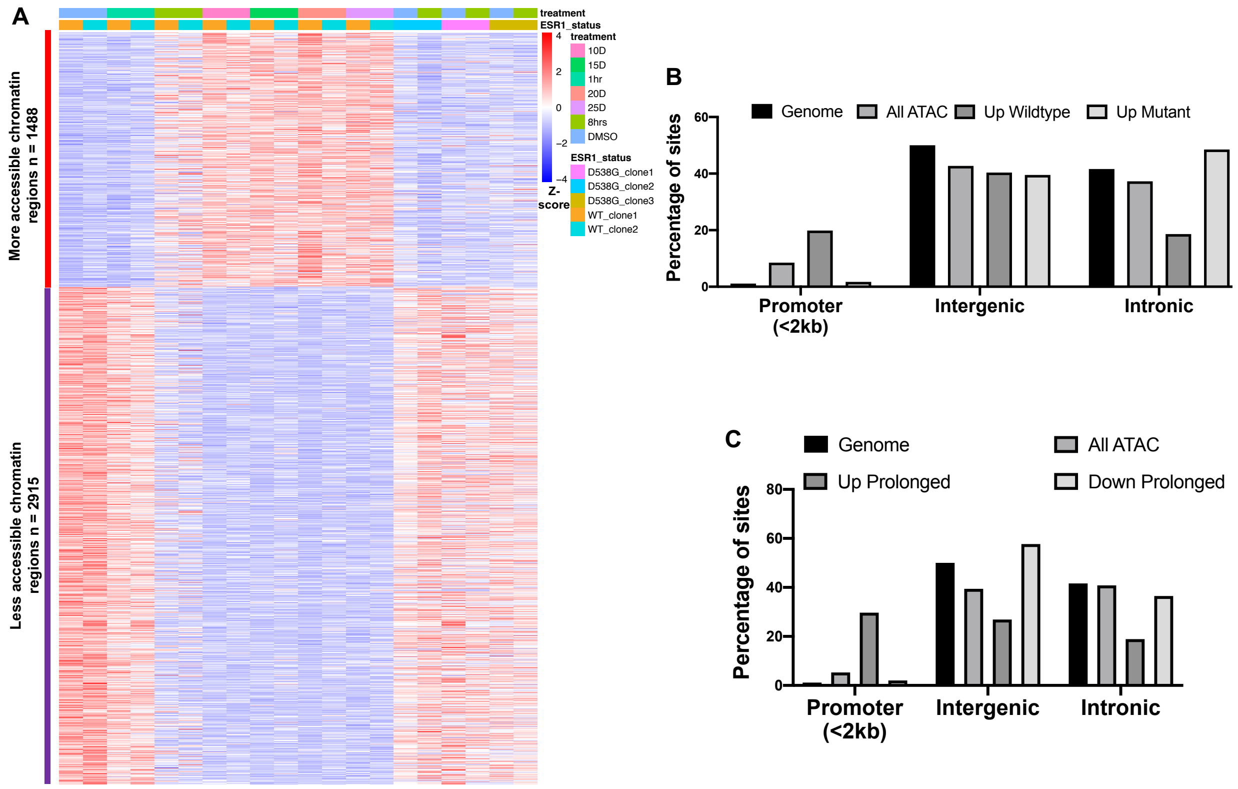


**Supplemental Figure 7. Prolonged ATAC-seq and ATAC-seq CEAS analysis (related to Figure 6) A.** Heatmap displays signal at ATAC-seq regions that change chromatin accessibility in wildtype ESR1 cells in response to prolonged E2 treatment. Sample types are indicated by the column annotations described in the legend. **B.** Genomic distribution of promoters, intergenic and intronic regions in all ATAC-seq regions of open chromatin, regions that are more accessible in wildtype ESR1 and D538G mutant cell lines. **C.** Genomic distribution of promoters, intergenic and intronic regions in all ATAC-seq regions of open chromatin and regions that are more accessible in wildtype ESR1 cell lines in response to the prolonged E2 induction.

**Supplemental Table 3**

Table S3, Related to Methods – Mutant ESR1 generation: Primers used to PCR amplify *ESR1* homology arms, gBlock used for template amplification and guide RNA sequences.

| Name | Sequence |
|---|---|
| ER_Fetch_arm1_F (homology arm upstream) | TCCCCGACCTGCAGCCCAGCT CTTGAACTGCTTTACTCA |
| ER_Fetch_arm1_R (homology arm upstream) | CCGGAACCTCCTCCGCTCCC GACCGTGGCAGGGAAACC |
| ER_Fetch_arm2_F (homology arm downstream) | AGTTCTTCTGATTCGAACATC CCACACGGTTCAGATAATCC |
| ER_Fetch_arm2_R (homology arm downstream) | TGGAGAGGACTTTCCAAG AACTTGCTGCTCTCCATAGG |
| gBlock (IDT) with D538G mutation in red | CTTGAACTGCTTTACTCATTTAAAATACCCACTCCTGCTTG GCTGAATATCTCATGTTGTCTTTTTAGAAGCTTTGGCGATC CTATTTGAATGCATTTAGGTCCTATTGGAGGGGAATAGGA TCTCATTTGAGGCCACGGAGGTCCATGGAAGTCACCTGC ATAGCAAATACCCTGAAAGTGGCTGCAGGGAGAGTGTGA GGGTGGGACCGCCCTGGTAGGAGGTGGAAAATGAAAAAC ACACGGCCATGAGTTCCAGATTAGGGCTTCTGAAAGCCC TCAGCTTTCCCAGCTCCCATCCTAAAGTGGGTCTTTAAAC AGGAAGAAAGAAAGATTGCTAAGTGTCTTTGGAGTTCCTC TTCCTTCCCCTTCTAGGGATTTCAGCACTCCTGGGGCTCG GGTTGGCTCTAAAGTAGTCCTTTCTGTGTCTTCCCACCTAC AGTAACAAAGGCATGGAGCATCTGTACAGCATGAAGTGCA AGAACGTGGTGCCCCTCTATGGCCTGCTGCTGGAGATGCT GGACGCCCACCGCCTACATGCGCCCACTAGCCGTGGAGG GGCATCCGTGGAGGAGACGGACCAAAGCCACTTGGCCAC TGCGGGCTCTACTTCATCGCATTCCTTGCAAAAGTATTACA TCACGGGGGAGGCAGAGGGTTTCCCTGCCACGGTCTGAG AGCTCCCTGGCTCCCACACGGTTCAGATAATCCCTGCTGCA TTTTACCCTCATCATGCACCACTTTAGCCAAATTCTGTCTCC TGCATACACTCCGGCATGCATCCAACACCAATGGCTTTCTA GATGAGTGGCCATTCATTTGCTTGCTCAGTTCTTAGTGGCA CATCTTCTGTCTTCTGTTGGGAACAGCCAAAGGGATTCCAA GGCTAAATCTTTGTAACAGCTCTCTTTCCCCCTTGCTATGTT ACTAAGCGTGAGGATTCCCGTAGCTCTTCACAGCTGAACTC AGTCTATGGGTTGGGGCTCAGATAACTCTGTGC |
| ER_stop_spCas9_gRNA1_top | CACCG CCACGGTCTGAGAGCTCCC |
| ER_stop_spCas9_gRNA1_bot_R2 | AAAC GGGAGCTCTCAGACCGTGG C |
| ER_stop_spCas9_gRNA2_top | CACCG CTGAACCGTGTGGGAGCCA |
| ER_stop_spCas9_gRNA2_bot_R2 | AAAC TGGCTCCCACACGGTTCAG C |

**Supplemental Table 4**

Table S4, Related to Methods – Primers to genotype *ESR1*

| Name | Sequence |
|---|---|
| ESR1-LBD_F4 (Sanger sequencing) | GGAAAGGCATTTAGATCGTATTCTGAG |
| ESR1-LBD_R1 (Sanger sequencing) | ATGAAGTAGAGCCCGCAGTG |
| ESR1-LBD_F4 (Tagged Allele) | GGAAAGGCATTTAGATCGTATTCTGAG |
| ESR1-LBD_STOP_R1 (Tagged Allele) | TGGTGCATGATGAGGGTAAA |

**Supplemental Table 5**

Table S5, Sanger sequencing results

| Clone name | Sanger sequencing results of *ESR1* mutation region |
|---|---|
| WT Clone 1 | TGANGTGAGAGANTTAACANTGGAGCGTCTTGAACTGCTTTACTCA<br>TTTAAAATACCCACTCCTGCTTGGCTGAATATCTCATGTTGTCTTTT<br>TAGAAGCTTTGGCGATCCTATTTGAATGCATTTAGGTCCTATTGGA<br>GGGGAATAGGATCTCATTTGAGGCCACGGAGGTCCATGGAAGTC<br>ACCTGCATAGCAAATACCCTGAAAGTGGCTGCAGGGAGAGTGTGA<br>GGGTGGGACCGCCCTGGTAGGAGGTGGAAAATGAAAAACACACG<br>GCCATGAGTTCCAGATTAGGGCTTCTGAAAGCCCTCAGCTTTCCC<br>AGCTCCCATCCTAAAGTGGGTCTTTAAACAGGAAGAAAGAAAGATT<br>GCTAAGTGTCTTTGGAGTTCCTCTTCCTTCCCCTTCTAGGGATTTC<br>AGCACTCCTGGGGCTCGGGTTGGCTCTAAAGTAGTCCTTTCTGTG<br>TCTTCCCACCTACAGTAACAAAGGCATGGAGCATCTGTACAGCAT<br>GAAGTGCAAGAACGTGGTGCCCCTCTATGACCTGCTGCTGGAGAT<br>GCTGGACGCCCACCGCCTACATGCGCCCACTAGCCGTGGAGGGG<br>CATCCGTGGAGGAGACGGACCAAAGCCACTTGGCCACTGCNNNN<br>NNNNNNNTTCANANNNA |
| WT Clone 2 | TGNNGTGNGAGANTTAACAATGGAGCGTCTTGAACTGCTTTACTC<br>ATTTAAAATACCCACTCCTGCTTGGCTGAATATCTCATGTTGTCTTT<br>TTAGAAGCTTTGGCGATCCTATTTGAATGCATTTAGGTCCTATTGG<br>AGGGGAATAGGATCTCATTTGAGGCCACGGAGGTCCATGGAAGT<br>CACCTGCATAGCAAATACCCTGAAAGTGGCTGCAGGGAGAGTGTG<br>AGGGTGGGACCGCCCTGGTAGGAGGTGGAAAATGAAAAACACAC<br>GGCCATGAGTTCCAGATTAGGGCTTCTGAAAGCCCTCAGCTTTCC<br>CAGCTCCCATCCTAAAGTGGGTCTTTAAACAGGAAGAAAGAAAGA<br>TTGCTAAGTGTCTTTGGAGTTCCTCTTCCTTCCCCTTCTAGGGATT<br>TCAGCACTCCTGGGGCTCGGGTTGGCTCTAAAGTANTCCTTTCTG<br>TGTCTTCCCACCTACNGTAACAAAGGCATGGAGCATCTGTACAGC<br>ATGAAGTGCAANNNNNTGGTGCCCCTCNATGACNTGCTGCTGGAG<br>ATGCTGGACGCCNACCGNCTACNNNNNNCCTNANACCCGTGNAN<br>GGGCAACNGTGNAGGANACCNACCAAAACNAATTGNCCACNGCN<br>CGCCNNNACTTTC |
| D538G Clone 1 | CTTATTNANTCGACCTTGGGNTGATCAGTTACTTGGAGGACTTGG<br>GGAAGTACCAGGCACTGCGGGCTCTACTTCATAAGTTTGGCGATC<br>CTAATTGAATGCATTCCCGTCCTATTGGAGGGTAATAAGATCTCAT<br>TTGAGGCCACAGAGGTCCATGGAACTCACCTGCATAATAAATACC<br>CTGAAATTGGTTGCAGGAATAGTGTAAGGGTGGGACCGCCCTGGT |

| | |
|---|---|
| | AGGAGGTGGAAAATGAAAAACGCACGGACATGAGTTCCAGATTAG GGCTTCTGAAAGCCCTCGTGTTTCCCAGCTCCCATCCTAGAGTGG GTCTTTACACCTGAATANAGAAAGATTGCTAAGTGTCTTTGGAGTT CCTCTTCCTTCCCCTTCTAGGGATTTCAGCACTCCTGGGGCTCGG GTTGGCTCTAAAGTAGTCCTTTCTGTGTCTTCCCACCTACAGTAAC AAAGGCATGGAGCATCTGTACAGCATGAAGTGCAAGAACGTGGTG CCCCTCTATG(A*)CCTGCTGCTGGAGATGCTGGACGCCCACCGCC TACATGCGCCCACTAGCCGTGGAGGGGCATCCGTGGAGGAGACG GACCAAAGCCACTTGGCCACTGCNNNNNNTACTTNA |
| D538G Clone 2 | TGCNNTGNAGCGTCTTGAACTGCTTTACTCATTTAAAATAACCACT CCTGCTTGGCTGAATATCTCNTGTTGTCTTTTTACTAGCTTTGGCG ATCCTATTTGAATGCATTTAGGTCCTATTGGAGGGGAATAGGATCT CATTTGAGGCCACAGAGGTCCATGGAAGTCACCTGCATAACAAAT ACCCTGAAAGTGGCTGCAGGGAGAGTGTGAGGGTGGGACCGCCC TGGTAGGAGGTGGAAAATGAAAAACACACGGCCATGAGTTCCAGA TTAGGGCTTCTGAAAGCCCTCAGCTTTCCCAGCTCCCATCCTAGA GTGGGTCTTTANACAGGAAAAAAGAAAGATTGCTAAGTGTCTTTGG AGTTCCTCTTCCTTCCCCTTCTAGGGATTTCAGCACTCCTGGGGCT CGGGTTGGCTCTAAAGTAGTCCTTTCTGTGTCTTCCCACCTACAGT AACAAAGGCATGGAGCATCTGTACAGCATGAAGTGCAAGAACGTG GTGCCCCTCTATG(A*)CCTGCTGCTGGAGATGCTGGACGCCCACC GCCTACATGCGCCCACTAGCCGTGGAGGGGCATCCGTGGAGGAG ACGGACCAAAGCCACTTGGCCACTGNNNNNNNNNNNNTTNNNANN NNNNNNNNCCGNTNNNNNNNGNCNGNGNNANNGNNANGTGNNN AAAGANNNAANNGNCNNNCTTNANNGNCNNNNTCGANNNNNTGG AGNGGNNNNNGNCNNTNNNANNNG |
| D538G Clone 3 | TNCANTGGAGCGTCTTGAACTGCTTTACTCATTTAAAATACCCACT CCTGCTTGGCTGAATATCTCNTGTTGTCTTTTTACTANCTTTGGCG ATCCTATTTGAATGCATTTAGGTCCTATTGGAGGGGAATAGGATCT CATTTGAGGCCACGGAGGTCCATGGAAGTCACCTGCATAGCAAAT ACCCTGAAAGTGGCTGCAGGGAGAGTGTGAGGGTGGGACCGCCC TGGTAGGAGGTGGAAAATGAAAAACACACGGCCATGAGTTCCAGA TTAGGGCTTCTGAAAGCCCTCAGCTTTCCCAGCTCCCATCCTAAA GTGGGTCTTTAAACAGGAATAAAGAAAGATTGCTAAGTGTCTTTGG AGTTCCTCTTCCTTCCCCTTCTAGGGATTTCAGCACTCCTGGGGCT CGGGTTGGCTCTAAAGTAGTCCTTTCTGTGTCTTCCCACCTACAGT AACAAAGGCATGGAGCATCTGTACAGCATGAAGTGCAAGAACGTG GTGCCCCTCTATG(A*)CCTGCTGCTGGAGATGCTGGACGCCCACC GCCTACATGCGCCCACTAGCCGTGGAGGGGCATCCGTGGAGGAG ACGGACCAAAGCCACTTGGCCACTGCGGNNNNTACTTCA |

*Location of Heterozygous A/G Variant

**Supplemental Table 6**

Table S6, Related to Methods – qPCR primer sequences

| Name | Sequence |
|---|---|
| PGR_qPCR_F | ACCCGCCCTATCTCAACTACC |
| PGR_qPCR_R | AGGACACCATAATGACAGCCT |
| MMP17_qPCR_F | CACTCATGTACTACGCCCTCA |
| MMP17_qPCR_R | TGGAGAAGTCGATCTGGATGTC |
| EHF_qPCR_F | CAGTGCAGTAGTGACCTGTTC |
| EHF_qPCR_R | CTGTGCTACCATAGTTGGTGTC |
| EPHA3_qPCR_F | CTGCTCTGTTCTCGACAGCTT |
| EPHA3_qPCR_R | CAGCTCCCCTTGAATTGTTTTTG |
| CTCF_qPCR_F | ACCTGTTCCTGTGACTGTACC |
| CTCF_qPCR_R | ATGGGTTCACTTTCCGCAAGG |
